# Top2a promotes the development of social behavior via PRC2 and H3K27me3

**DOI:** 10.1101/2021.09.20.461107

**Authors:** Yijie Geng, Tejia Zhang, Sean C. Godar, Brock R. Pluimer, Devin L. Harrison, Anjali K. Nath, Jing-Ruey Joanna Yeh, Iain A. Drummond, Marco Bortolato, Randall T. Peterson

## Abstract

Human infants exhibit innate social behaviors at birth, yet little is understood about the embryonic development of sociality. We screened 1120 known drugs and found that embryonic inhibition of topoisomerase IIα (Top2a) resulted in lasting social deficits in zebrafish. In mice, prenatal Top2 inhibition caused behavioral defects related to core symptoms of autism, including impairments in social interaction and communication. Mutation of Top2a in zebrafish caused downregulation of a set of genes highly enriched for genes associated with autism in humans. Both the Top2a-regulated and autism-associated gene sets possess binding sites for polycomb repressive complex 2 (PRC2), a regulatory complex responsible for H3K27 trimethylation. Moreover, both gene sets are highly enriched for H3K27me3. Inhibition of PRC2 component Ezh2 rescued social deficits caused by Top2 inhibition. Therefore, Top2a is a key component of an evolutionarily conserved pathway that promotes the development of social behavior through PRC2 and H3K27me3.

## INTRODUCTION

Sociality is broadly conserved across the animal kingdom, facilitating cooperation, reproduction, and protection from predation. Human infants exhibit an innate social preference behavior at birth: infants less than an hour old fix their gaze on human face-like images longer than other images^1^. Throughout life, this simple innate social drive serves as a foundation for the development of complex and versatile adult social behaviors. On the other hand, social dysfunction is a hallmark of several neurodevelopmental disorders^2^ such as autism spectrum disorder. Deficits in early social behavior are linked to later diagnoses of autism^3^. The onset of autism can be traced back to the prenatal stage^4-6^, which coincides with the development of innate social behavior.

Despite its importance, little is known about the embryonic development of sociality. While many genes conferring a small amount of autism risk have been identified by genome-wide association studies (GWAS)^7-10^, it is much less clear how they each contribute to autism etiology. Apart from these genetic factors, environmental factors are estimated to account for ∼40% of autism risk^11,12^, yet few have been identified. In this study, we investigated the development of sociality in zebrafish (*Danio rerio*) using a chemical genetics approach. The zebrafish has become an important model organism for behavioral studies due to its behavioral and neurophysiological similarities to humans^13-17^. It possesses an evolutionarily conserved social brain network^15^, enabling us to acquire insights into social behavior development of potential relevance for mammals^18-21^. Like in humans, a robust social preference behavior emerges early in zebrafish^22-25^. To restrict this investigation to developmental events occurring during the embryonic stage, we conducted a chemical screen that allows us to precisely apply and withdraw chemicals to and from zebrafish embryos during embryonic development. We screened an annotated library containing chemicals with known biological targets to help rapidly identify downstream biological targets.

Using an automated high-throughput behavioral analysis system which we named Fishbook, we screened 1120 known drugs and found that embryonic inhibition of topoisomerase II α (Top2a) resulted in social deficits in juvenile zebrafish. In mice, prenatal Top2 inhibition caused postnatal behavioral defects that are specifically related to the two core symptoms of autism: social communication impairments and increased perseveration. In fact, several known environmental risk factors of autism are Top2 inhibitors^26-29^, indicating that Top2 inhibition may be a common mechanism through which environmental insults contribute to autism risk.

To elucidate the Top2-mediated mechanism of action, we performed RNA-seq of Top2a mutant zebrafish and observed downregulation of a set of genes highly enriched for autism risk genes. Using a custom upstream analysis pipeline, we found that both the Top2a-dependent gene set and a set of autism-associated genes maintained by the Simons Foundation for Autism Research (SFARI genes hereafter) are selectively targeted by polycomb repressive complex 2 (PRC2), a regulatory complex responsible for depositing H3K27 trimethylation (H3K27me3). In corroboration of this finding, we found both gene sets to be highly enriched for H3K27me3. Strikingly, chemical inhibition of the PRC2 component Ezh2 rescued social deficits caused by Top2 inhibition in zebrafish, indicating that Top2a likely functions by antagonizing PRC2/H3K27me3-mediated gene silencing. These findings identify Top2a as a key component of an evolutionarily conserved pathway that promotes the development of social behavior through PRC2 and H3K27me3.

## RESULTS

### Fishbook: a scalable social behavior assay system

Zebrafish develop a robust social preference for age-matched conspecifics at 3 weeks of age^22^. To quantitatively assess this behavior, we developed a scalable and fully automated assay system named Fishbook. Briefly, a test arena (Figure 1A) was manufactured by 3D printing (Figure 1B). Each arena (Figure 1C, red rectangle) is a long rectangular lane printed using nontransparent material, divided into 3 parts by two transparent windows (4): the longer middle part serves as the test compartment (2), one of the two smaller end-compartments contains a live fish as social stimulus (1), and the other end-compartment is left empty (3) (Movies S1). To maximize throughput, we grouped 44 arenas together (Figure 1D), and constructed a unique telecentric lens-based imaging system which enables us to simultaneously image all 44 arenas while avoiding optical obstruction by the non-transparent arena walls (Figure 1E). The Fishbook system is capable of screening >1000 fish per day.

**Figure 1.**
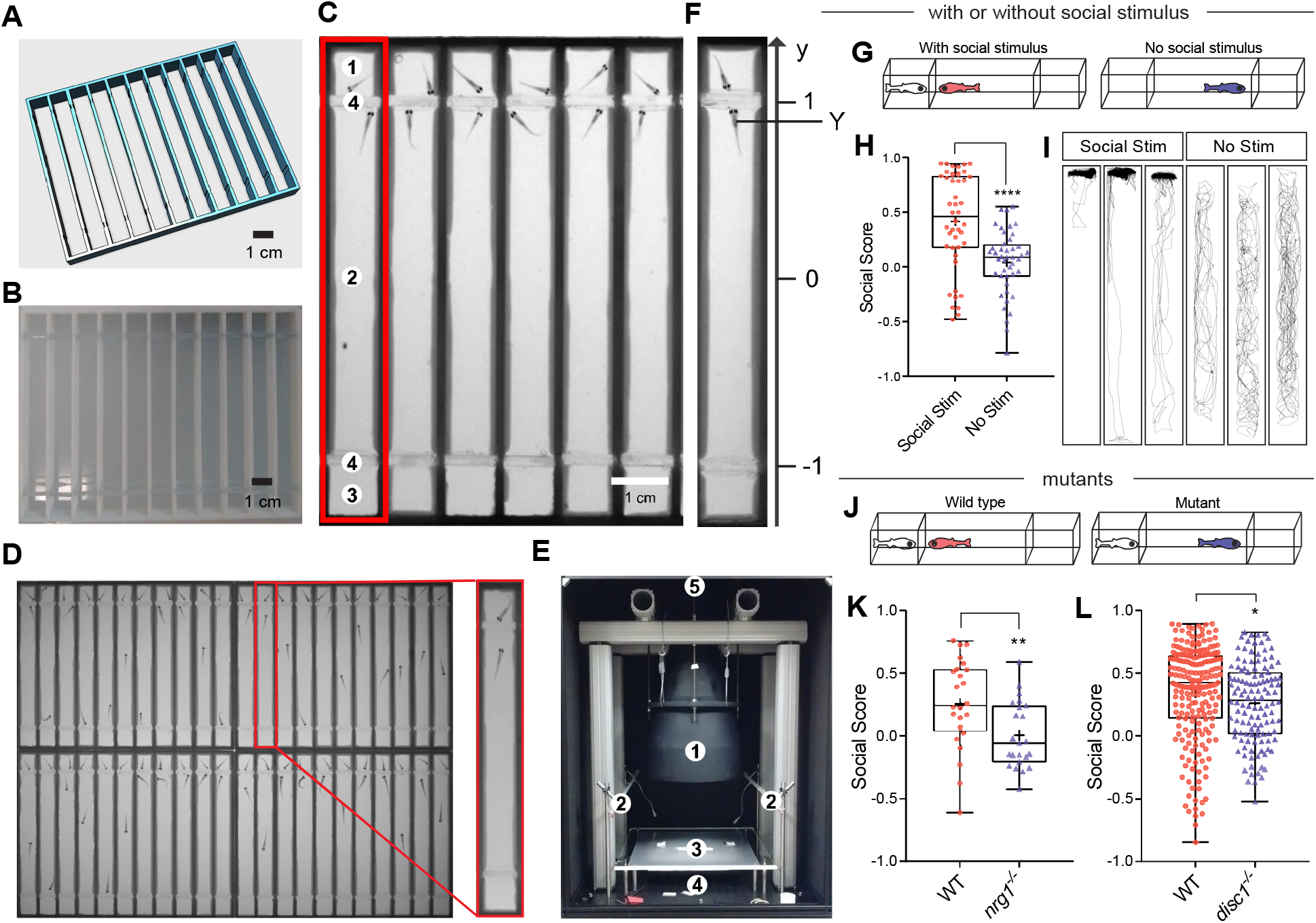
The Fishbook assay. (**A, B**) The design and a 3D printed array of Fishbook arenas. (**C**) Screen shot of a Fishbook assay. Red rectangle: a Fishbook arena; 1: social stimulus compartment; 2: social test compartment; 3: empty control compartment; 4: transparent windows. (**D**) An array of 44 arenas. (**E**) The Fishbook imaging station. 1: telecentric lens; 2: white light LED panels providing ambient light; 3: sample deck; 4: 850 nm infrared LED backlight panel; 5: CCD camera. (**F**) Social score is calculated as the average Y-axis position for the duration of a test. (**G-I**) Fishbook with or without social stimulus fish (G), showing boxplots (H) and representative tracking plots (I) of WT fish tested with social stimulus (Social Stim; n = 44) or without social stimulus (No Stim; n = 44). In each boxplot, box encloses data points from the 25^th^ percentile to the 75^th^ percentile, horizontal line and cross mark the median and the mean, the lines above and below the box reach datapoints with the maximum and minimum values. ****: *p*<0.0001. (**J-L**) Fishbook detects social deficits in mutants (J), comparing WT (n=24) and homozygous *nrg1* knockout (*nrg1*^*-/-*^; n=23) fish, both from a heterozygous (*nrg1*^*+/-*^) incross and genotyped individually after Fishbook assay (K), and WT sibling (n=209) and homozygous *disc1* knockout (*disc1*^*-/-*^; n=122) fish (L). *: *p*<0.05, **: *p*<0.01.

A social score with values ranging between −1 and 1 was defined to quantitatively measure social behavior (Figure 1F). Wild-type (WT) fish typically spend a significant amount of time near the social stimulus fish, generating high social scores; without a social stimulus, test subjects swim randomly, giving rise to an average social score close to 0 (Figure 1G-1I). Consistent with a previous report^22^, this preference is lost in the dark (Figure S1A), indicating that it is driven primarily by visual cues. Social score is also drastically reduced when social stimulus fish is replaced by a pebble of similar size (Figure S1B), suggesting that this preference behavior is highly specific to social cues and cannot be explained solely by curiosity toward a novel object. To validate its power in recognizing social deficits, we examined fish possessing loss-of-function mutations of genes linked to autism and schizophrenia, including *Nrg1*^30^ (Figures 1K & S1C) and *Disc1*^31^ (Figures 1L & S1D). Social scores are significantly reduced in both mutants, demonstrating that the Fishbook assay can effectively detect social defect caused by a single genetic mutation.

### A systematic screen identifies fluoroquinolones as inhibitors of social development

Using Fishbook, we screened the Prestwick library which contains 1120 compounds with known biological targets. Briefly, embryos were exposed to chemicals during the first 3 days of development (0 - 3 days post fertilization, or dpf), raised to 3 weeks of age, and tested in the Fishbook assay (Figure 2A). Hits were defined as having an average social score < 0.1. Due to the relatively large variation expected for a behavior-based assay, to reduce the number of false positive hits, primary screening was performed in duplicates and hits were confirmed by repeat testing. Four confirmed hits were found, including flumequine, lomefloxacin, ofloxacin, and oxolinic acid (Figures 2B & S2A). Importantly, these four compounds were the only validated hits from the screen, and all belonged to the same class of antibiotics named fluoroquinolones. A previous publication reported modulation of zebrafish shoaling behavior by long term exposure to a mixture of 3 fluoroquinolones (ofloxacin, ciprofloxacin, and enrofloxacin) and 3 tetracyclines, suggesting a potential link between fluoroquinolones and zebrafish social behavior which is in line with our observation, but did not examine the effect of fluoroquinolones alone without tetracyclines or provide insights on their molecular mechanisms^32^.

**Figure 2.**
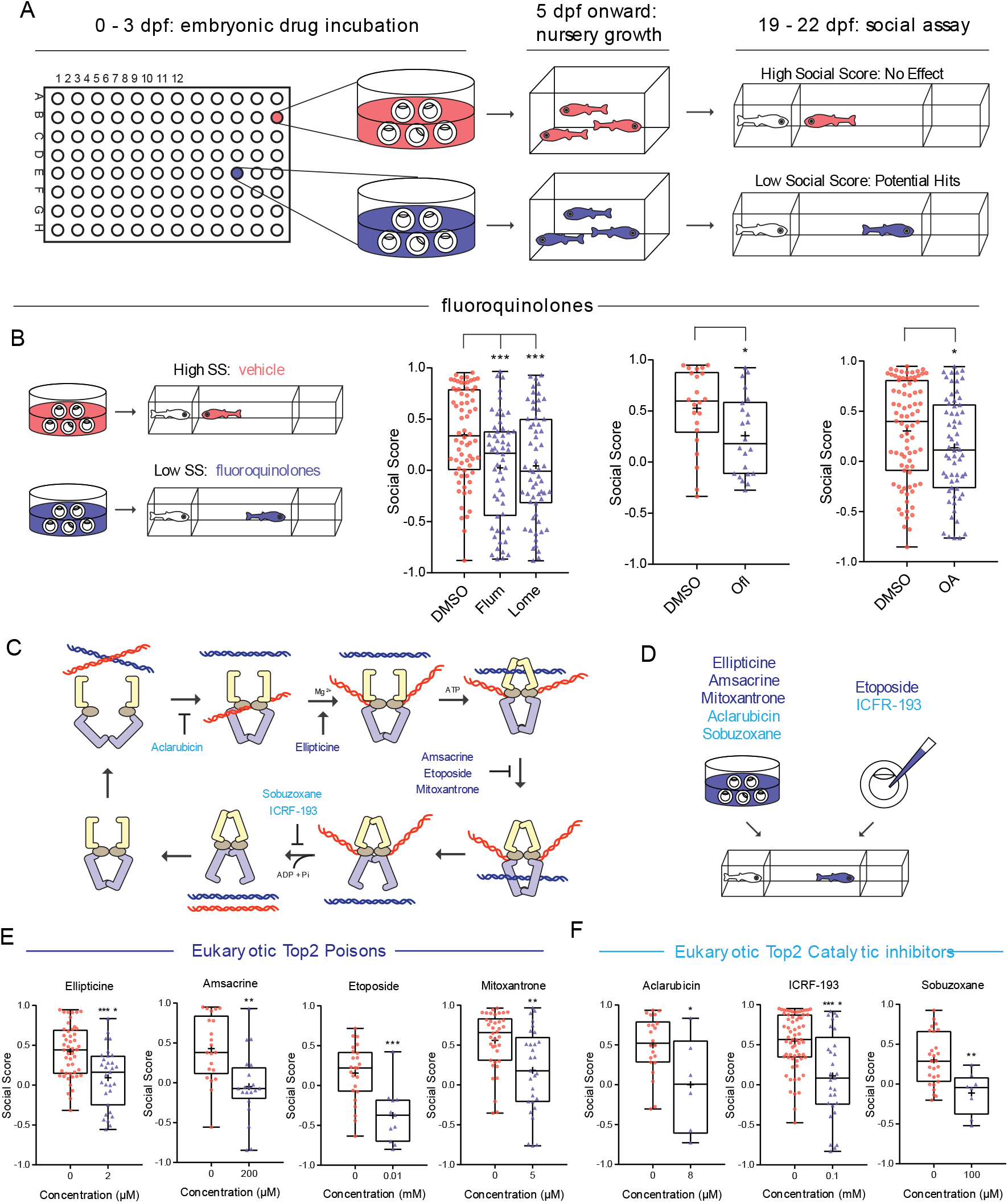
Screening for inhibitors of social development identifies fluoroquinolones and other Top2 inhibitors. (**A**) The screening workflow. (**B**) Fluoroquinolones induce social deficits. Comparing DMSO (n=68, 22, 82, from left to right) with flumequine (Flum, 25 μM, n=56), lomefloxacin (Lome, 100 μM, n=60), ofloxacin (Ofl, 200 μM, n=22), and oxolinic acid (OA, 200 μM, n=62). (**C**) The catalytic cycle of Top2 and inhibitors targeting different steps of the cycle. (**D-F**) Eukaryotic Top2 inhibitors induce social deficits (D). Comparing fish treated with vehicle control (0 μM in each panel), Top2 poisons ellipticine (0 μM: n=48; 2 μM: n=31), amsacrine (0 μM: n=21; 200 μM: n=20), etoposide (0 μM: n=24; 10 μM: n=11), and mitoxantrone (0 μM: n=38; 5 μM: n=29) (E), and Top2 catalytic inhibitors aclarubicin (0 μM: n=22; 8 μM: n=6), ICRF-193 (0 μM: n=65; 100 μM: n=29), and sobuzoxane (0 μM: n=27; 100 μM: n=8) (F). *: *p*<0.05, **: *p*<0.01, ***: *p*<0.001, ****: *p*<0.0001.

### Top2 inhibition is responsible for the social deficit phenotype

Although fluoroquinolones are best known as antibiotics targeting the prokaryotic type-II topoisomerases, they also inhibit the eukaryotic topoisomerase II (Top2), albeit at higher doses. For example, the IC_50_ of ofloxacin and lomefloxacin against HeLa cell Top2 are 365 μM and 613 μM^33^, and their effective doses in the Fishbook assay are 200 μM and 100 μM, respectively. Prior studies have also suggested a link between Top2 and autism^34-38^ (see Discussion for details). We thus hypothesized that fluoroquinolones inhibit the development of sociality by targeting zebrafish Top2.

To test this hypothesis, we examined several structurally diverse eukaryotic Top2 inhibitors (Figure S2B). They target different steps of the Top2 catalytic cycle^39,40^ (Figure 2C) and can be grouped into two categories based on their mechanisms of action, including Top2 poisons (ellipticine, amsacrine, etoposide, and mitoxantrone), and Top2 catalytic inhibitors (aclarubicin, ICRF-193, and sobuzoxane). Through embryonic exposure (Figure 2D), all 7 structurally diverse Top2 poisons and inhibitors induced social deficits (Figures 2E & 2F). We also tested several environmental chemicals which were known to inhibit eukaryotic Top2 activity, including chlorpyrifos, chlorpyrifos oxon, and genistein (see Discussion for more information), and observed similar social deficits (Figure S2C). These results demonstrated that inhibition of zebrafish Top2 can indeed induce social deficits.

### Top2 inhibition does not function by inhibiting transcription or genomic stability

Apart from its role in facilitating replication, Top2 plays important parts in promoting transcription and genomic stability^41^. We therefore investigated whether the effect of Top2 inhibition can be attributed to deficits in one of these two apparent mechanisms. We first examined two RNA polymerase II inhibitors, actinomycin D and triptolide, but did not observe social deficits (Figures S2D & 2E). We then tested two DNA double strand break (DSB) inducers, bleomycin and hydroxyurea, and again found that neither compound induced social deficits (Figure S2F). This result is also supported by the fact that while Top2 poisons induce DSBs, catalytic inhibitors do not, yet both classes of Top2 inhibitors cause social deficits (Figures 2E & 2F). Together, these data support the possibility that Top2 inhibition functions through a mechanism other than general transcription inhibition or DNA instability.

### Top2a, but not Top2b, is required for the development of social behavior

We next investigated which isoform, Top2a or Top2b, was responsible for the social deficit phenotype. We first applied two splice-blocking morpholino oligonucleotides (MO) (Figure S3A). Compared to permanent genomic editing methods such as CRISPR, MO’s transient-acting nature better mimics effects of the Top2 inhibitors. Note that maternal Top2a mRNAs are present during the first 24 hours of embryogenesis^42^ and will escape inhibition by Top2a-MO. Following injection, Top2a-MO induced social deficits, which were rescued by co-injecting *hTOP2A* mRNA (Figures 3A & S3B). Top2b-MO, however, failed to induce social deficits (Figure 3B). A Top2a-selective inhibitor sodium salicylate also induced social deficits (Figure 3C) when applied at a dose known to confer its isoform selectivity^40^.

**Figure 3.**
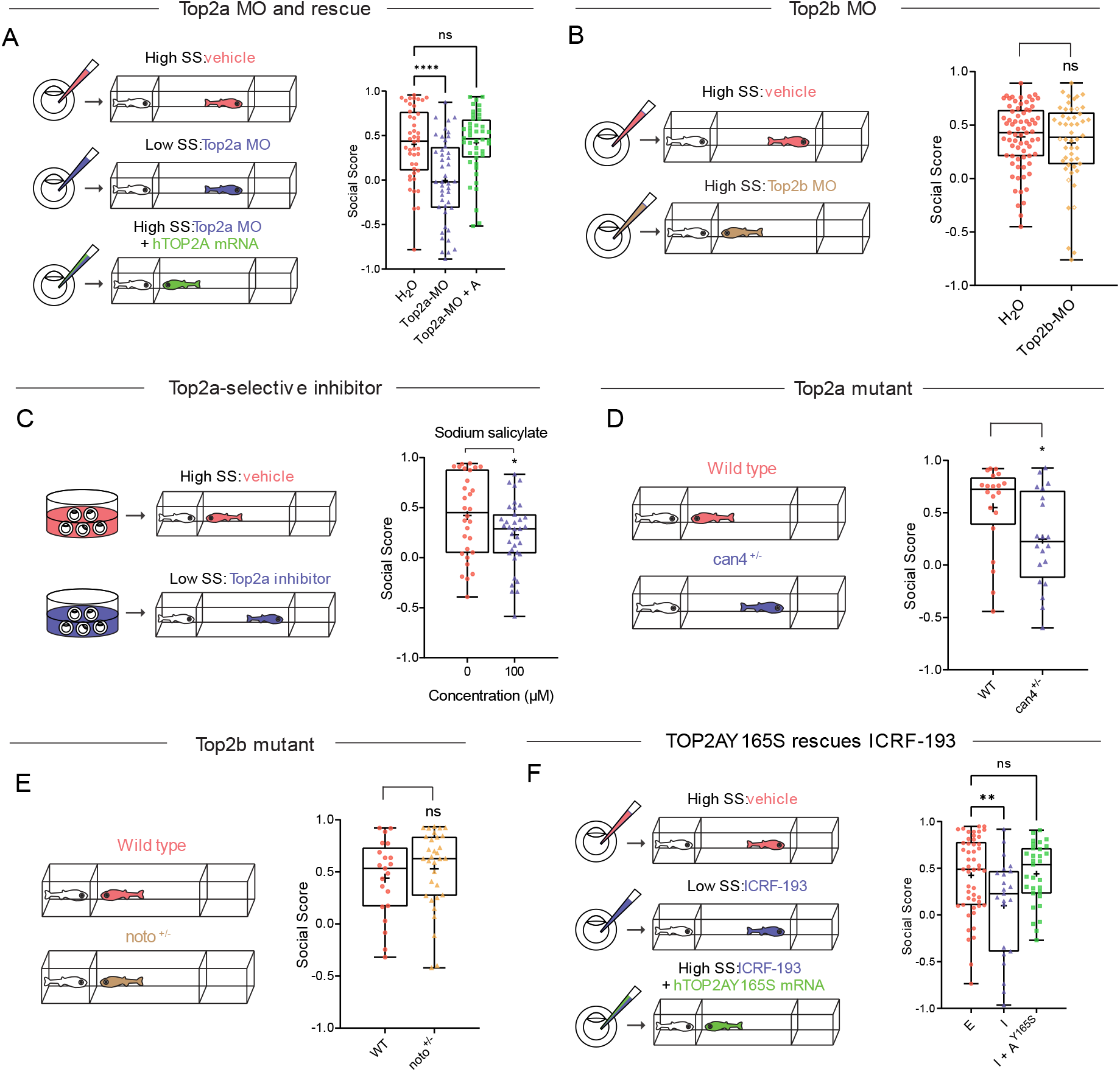
Top2a rather than Top2b is required for the development of social behavior. (**A**) Top2a-MO (0.05 mM, n=44) induces social deficits compared to vehicle (H_2_O, n=44), which is rescued by *hTOP2A* mRNA (250 ng/μl, Top2a-MO+A, n=44). (**B**) Top2b-MO (0.05 mM, n=50) does not induce social deficits compared to vehicle (H_2_O, n=71). (**C**) Top2a selective inhibitor sodium salicylate (100 μM) induces social deficits. (**D**) Heterozygous *can4* mutant (*can4*^*+/-*^, n=20) exhibits social deficits compared to WT (n=20). Both mutant and WT were acquired from a heterozygous (*can4*^*+/-*^) incross and genotyped individually after Fishbook assay. (**E**) Heterozygous *noto* mutant (*noto*^*+/-*^, n=32) shows no social deficits compared to WT (n=21). Both mutant and WT were acquired from a heterozygous (*noto*^*+/-*^) incross and genotyped individually after Fishbook assay. (**F**) ICRF-193 (I, 0.1 mM, n=23) induced social deficits compared to the vehicle control ethanol (E, 1%, n=50), which is rescued by *hTOP2AY165S* mRNA (250 ng/μl, I+A^Y165S^, n=31). ns: not significant, *: *p*<0.05, **: *p*<0.01, ****: *p*<0.0001.

We then examined two loss-of-function (LoF) mutants of Top2a (*can4*^43^) and Top2b (*noto*^44^). Because homozygous mutants are lethal, we established breeding colonies using heterozygous (het) mutants. Incrossing het mutants generated mixed populations composed of ∼ 1/3 WT and 2/3 het offspring. They were assayed blind in Fishbook, i.e., with no prior knowledge of each fish’s genetic background, and genotyped individually. Het Top2a mutant (*can4*^*+/-*^) exhibited reduced social score compared to WT sibling (Figure 3D), while het Top2b mutant (*noto*^*+/-*^) did not show reduction in social score (Figure 3E).

Finally, we tested whether transient overexpression of Top2a can rescue the Top2 inhibitor-induced social deficit. A potential concern is that overexpressed WT hTOP2A protein may simply bind to and “soak up” a Top2 inhibitor, leaving less free compound to interact with its real target, resulting in false positive rescue. We therefore generated a hTOP2A mutant (Y165S) which is functionally active but resistant to ICRF-193^45^. Transient overexpression of hTOP2AY165S by mRNA injection successfully rescued ICRF-193 (Figures 4L & S3C), demonstrating that Top2a alone can rescue Top2 inhibition. Together, these results suggest that Top2a is the isoform responsible for the social deficit phenotype.

**Figure 4.**
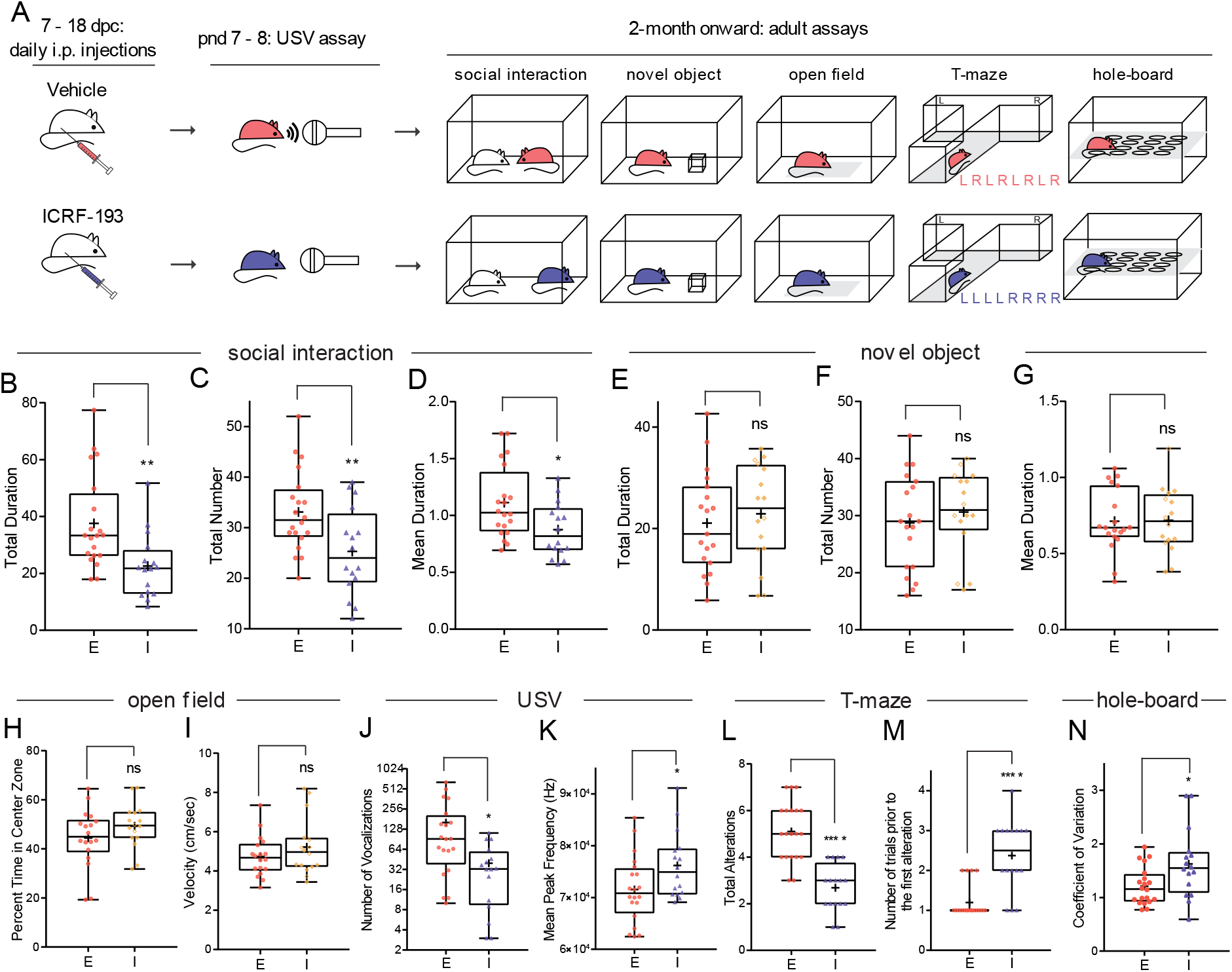
Prenatal inhibition of Top2 induces social communication impairments and increased perseveration in mouse. (**A**) Experimental design for prenatal ICRF-193 injection and subsequent experiments for the offspring. (**B-D**) ICRF-193 (I, n=16) reduces total duration (B), total number (C), and mean duration (D) of social investigation compared to vehicle control ethanol (E, n=20). (**E-G**) ICRF-193 (n=16) does not reduce total duration (E), total number (F), and mean duration (G) of novel object investigation compared to ethanol (n=19). (**H, I**) ICRF-193 (n=16) does not change percentage of time spent in the center zone (H) and velocity (I) of mice in open field assay compared to ethanol (n=18). (**J, K**) ICRF-193 (n=16) reduces total number (J) but increases mean peak frequency (K) of ultrasonic vocalization compared to ethanol (n=20). (**L, M**) ICRF-193 (n=16) reduces total number of alterations (number of times a mouse enters an alternative arm compared to the previous trial) (L) while increases number of trials prior to the first alteration (M) compared to ethanol (n=20) during a T-maze assay consisted of 8 trials. (**N**) ICRF-193 (n=16) increases coefficient of variation in hole-board assay compared to ethanol (n=20). ns: not significant, *: *p*<0.05, **: *p*<0.01, ***: *p*<0.001, ****: *p*<0.0001.

### Prenatal inhibition of Top2 induces social deficits and other autism-related behavioral defects in mice

To determine whether this Top2-mediated mechanism is conserved in mammals, we assessed the effect of prenatal Top2 inhibition in mice. Pregnant mice received daily intraperitoneal (i.p.) injections of ICRF-193 or vehicle control from 7 to 18 days post coitum (dpc). Offspring grew to adult stage without observable differences in gross morphology or body weight (Figures S3D) compared to vehicle control or no-treatment groups. Behavioral assays were conducted 2 – 10 months after birth (Figure 4A).

We first conducted a reciprocal social interaction assay^46^, in which 2 sex and age matched mice, including a test subject and a social stimulus mouse, were placed into a novel cage and video recorded. The number and duration of interactions between a test subject and the control mouse were manually scored by a blinded reviewer. ICRF-193-treated mice showed significantly reduced total duration, total number, and mean duration (Figures 4B-4D) of social investigations compared to the vehicle control. We then performed a novel object exploration assay to determine if the ICRF-193 mice lack interest in or even avoid novel stimuli in general. ICRF-193 mice did not show significantly altered object exploration compared to controls, as quantified by the total duration, total number, and mean duration (Figures 4E-4G) of their interactions with the novel object (a wooden cube). We also observed no significant changes in motion patterns in an open field assay, as measured by the time spent in the center of the test arena and average velocity (Figures 4H-4I), indicating intact motor capabilities and normal stress responses for the ICRF-193 mice. These results indicate that social behaviors are selectively targeted by prenatal Top2 inhibition, while exploratory behaviors, motor activity, and stress response remain intact.

We then investigated other autism-related behavioral deficits including impaired communication and perseveration. We examined ultrasonic vocalization (USV) of the pups for potential social communication deficits. When briefly separated from their mothers, ICRF-193-treated pups exhibited significantly reduced numbers of USVs compared to controls (Figure 4J), indicating a significant impairment in their communication. The ICRF-193-treated pups also showed a significantly elevated mean peak frequency in their vocalizations (Figure 4K), a phenotype that was also found in established mouse models of autism^47,48^. We then examined restrictive and repetitive behaviors using T-maze^46^ and hole-board^46^ assays. ICRF-193 mice repeatedly explored the same arm of a T-maze, as shown by their significantly reduced number of alternative explorations and delayed onset of first alteration (Figures 4L-4M) compared to the controls. Hole-board assay also revealed significantly elevated coefficient of exploratory variation, a measure of repetitive tendencies^46^, in ICRF-193 mice (Figure 4N). These results demonstrate that prenatal inhibition of Top2 induces behavioral deficits related to the core symptoms of autism in mice.

### Top2a depletion selectively downregulates autism risk genes

Based on the observed link between Top2a inhibition and autism-related behavioral deficits, we investigated whether Top2a depletion can selectively modulate expression of autism risk genes using RNA-seq of 3 dpf *can4*^*-/-*^ (Top2a) mutants (Figure 5A). Due to the presence of maternal Top2a mRNA during the first 24 hours of embryogenesis^42^, *can4*^*-/-*^ embryos are expected to be partially depleted of Top2a during the first few days of embryonic development. We found 8634 significantly (adjusted *p*-value < 0.05) upregulated genes and 8589 significantly downregulated genes in *can4*^*-/-*^ mutants compared to WT controls. Top2a was downregulated as expected (adjusted *p*-value = 2.16E-73). We used the human orthologs of zebrafish genes for subsequent analyses, which include 5044 human orthologs for the downregulated genes (hereafter referred to as *can4*^*-/-*^ downregulated genes, or can4Dn) and 5054 human orthologs for the upregulated genes (*can4*^*-/-*^ upregulated genes, or can4Up) (Data S1).

**Figure 5.**
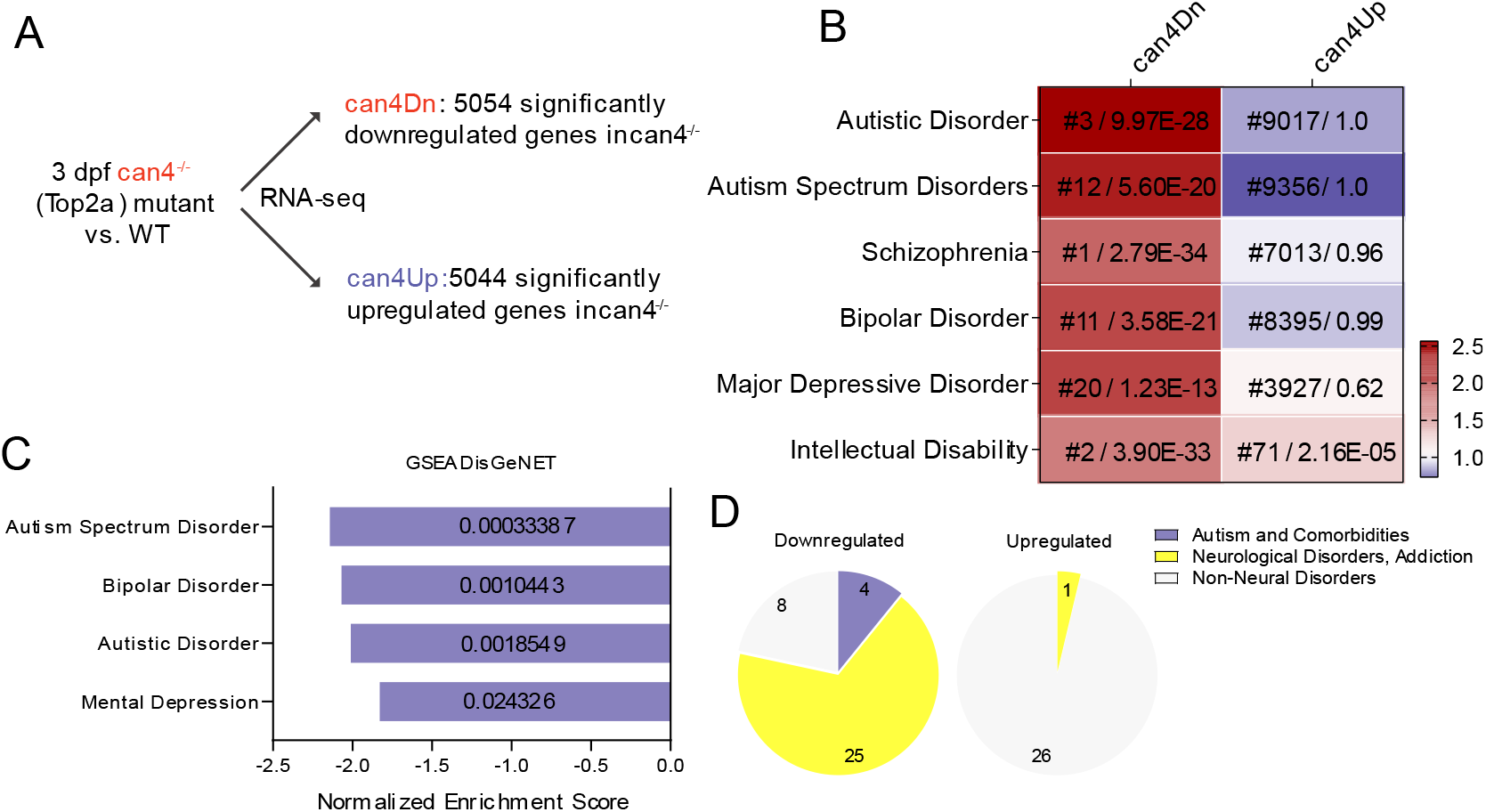
RNA-seq reveals downregulation of autism risk genes caused by Top2a depletion. (**A**) RNA-seq experimental design. (**B**) can4Dn but not can4Up is selectively enriched for autism and its comorbidities risk genes. For each cell, color represents odds ratio, and numbers represent ranking out of 24166 diseases in DisGeNET library (before slash) and adjusted *p*-value (after slash). (**C, D**) GSEA analysis using the DisGeNET library shows enrichment for autism and its comorbidities risk genes (C) and neurological conditions risk genes (D) in downregulated. Value inside each bar represents FDR (C). Significance: FDR<5%.

To identify diseases that most closely associate with genes affected by Top2a depletion, we analyzed two disease-gene libraries DisGeNET^49^ and GLAD4U^50^ using over-representation analysis (ORA) to search for disease risk gene sets that are enriched (over-represented) in the can4Dn and can4Up gene lists. can4Dn is enriched for genes associated with autism and its comorbid conditions including schizophrenia, bipolar disorder, depression, and intellectual disability; these diseases consistently rank at the top of the list among the 24166 diseases in the DisGeNET library and the 3071 in GLAD4U, demonstrating a high degree of specificity for these diseases (Figures 5B & S4A). can4Up is not enriched for most of these diseases except modestly for intellectual disability in DisGeNET. We also found a similar pattern of enrichment in independently curated disease gene sets related to autism and its comorbidities (SFARI^51,52^, AutismKB 2.0^53^, DisGeNET^49^, GLAD4U^50^, BDgene^54^, PsyGeNET^55^, SZGene^56^, SysID^57^, and GEPAD^58^) (Figure S4B). Using gene set enrichment analysis (GSEA) as an orthogonal approach, we again found significant enrichment (5% FDR) of autism and its comorbidities risk genes only in the downregulated genes (Figures 5C, 5D, S4C, & S5).

Pathway (KEGG and REACTOME) and gene ontology (GO) ORA analyses also revealed enrichment of genes related to pathways that are believed to contribute to autism etiology^59-62^, including axon guidance, neuronal development, glutamatergic signaling, and synaptic transmission, in can4Dn but not can4Up (Figure S6A). GSEA analysis using these libraries showed similar enrichments only in the downregulated genes (Figure S6B).

To eliminate any potential bias in our enrichment analyses caused by naturally-occurring enrichment or depletion of autism-associated genes in zebrafish, we examined the specificity of our analyses using a permutation assay. Because can4Dn and can4Up both contain close to 5000 genes, we randomly selected 1000 lists of 5000 genes out of the 14989 human genes with zebrafish orthologs and repeated the ORA analyses for these gene sets to create a null distribution. We found that can4Dn but not can4Up consistently gives significantly higher odds ratios compared to the null distribution, thus validating the specificity of the observed enrichment of autism risk genes (Figures S7-S9).

### can4Dn and autism risk genes share similar upstream regulators

We analyzed a published human TOP2A ChIP-seq dataset to investigate whether TOP2A selectively binds to autism risk genes^63^. We did not find enrichment of autism risk genes in TOP2A target genes (Figure 7B, first column), consistent with the traditional view of Top2a’s role as a “housekeeping” gene with no known selectivity for autism risk genes. Given that Top2a binding is not enriched in autism risk genes but Top2a-dependent regulation is, we hypothesized that some specificity factor may associate with Top2a, enabling its influence to be heightened in autism risk genes. To identify such a specificity factor, we conducted ORA analyses to identify upstream regulators that are significantly enriched in the promoter regions of can4Dn genes but not can4Up genes. We also analyzed SFARI genes in parallel as a representative set of autism risk genes. Strikingly, by analyzing a curated library based on published ChIP-seq data (ChIP-X^64,65^), we found that the SFARI genes and can4Dn are targeted by a highly similar set of upstream regulators, especially the transcription factor SUZ12 and its associating partner EZH2 (Figure 6A).

**Figure 6.**
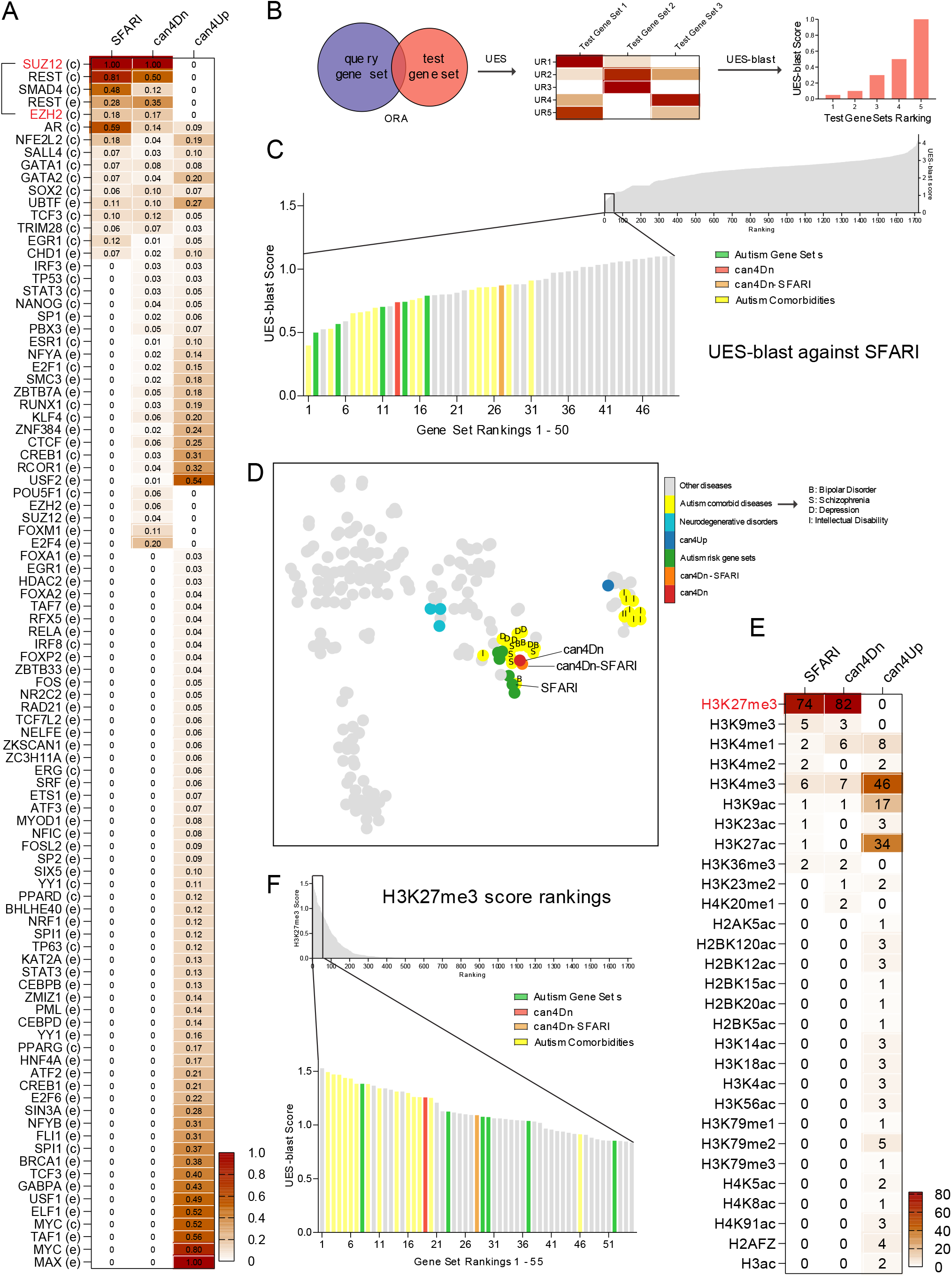
Upstream analyses find PRC2 and H3K27me3 as common upstream regulators of can4Dn and autism risk genes. (**A**) SFARI genes and can4Dn, but not can4Up, share very similar enrichment profiles for upstream regulators. Value in each cell represents the significance score (Methods). In row labels, c: ChEA, e: ENCODE, indicating source of ChIP-seq data. (**B**) UES analysis and UES-blast workflow. ORA analysis finds significantly enriched upstream regulators for a test gene set using reference gene sets from the ChIP-X library. A significance score is calculated for each upstream regulator (Methods), creating a signature for each test gene set which we named Upstream Enrichment Signature (UES). A blast-style querying algorithm (UES-blast) then identifies test gene sets with the most similar UES compared to a query gene set, e.g., SFARI genes. UR: upstream regulator. (**C**) UES-blast (Methods) shows that can4Dn, can4Dn-SFARI, and several independently curated autism risk gene sets rank high among control gene sets against the query gene set SFARI genes, indicating that they share a very similar UES with the SFARI genes. (**D**) tSNE clustering of can4Dn, can4Dn-SFARI, autism risk gene sets, and DisGeNET gene sets (only those containing > 500 genes). Letters label autism comorbidities risk gene sets, I: intellectual disability; D: depression; S: schizophrenia; B: bipolar disorder. (**E**) Genes in the SFARI and can4Dn gene sets, but not in can4Up, are specifically enriched for the H3K27me3 mark. (**F**) can4Dn, can4Dn-SFARI, and autism risk gene sets possess the highest H3K27me3 scores when compared to the control gene sets.

**Figure 7.**
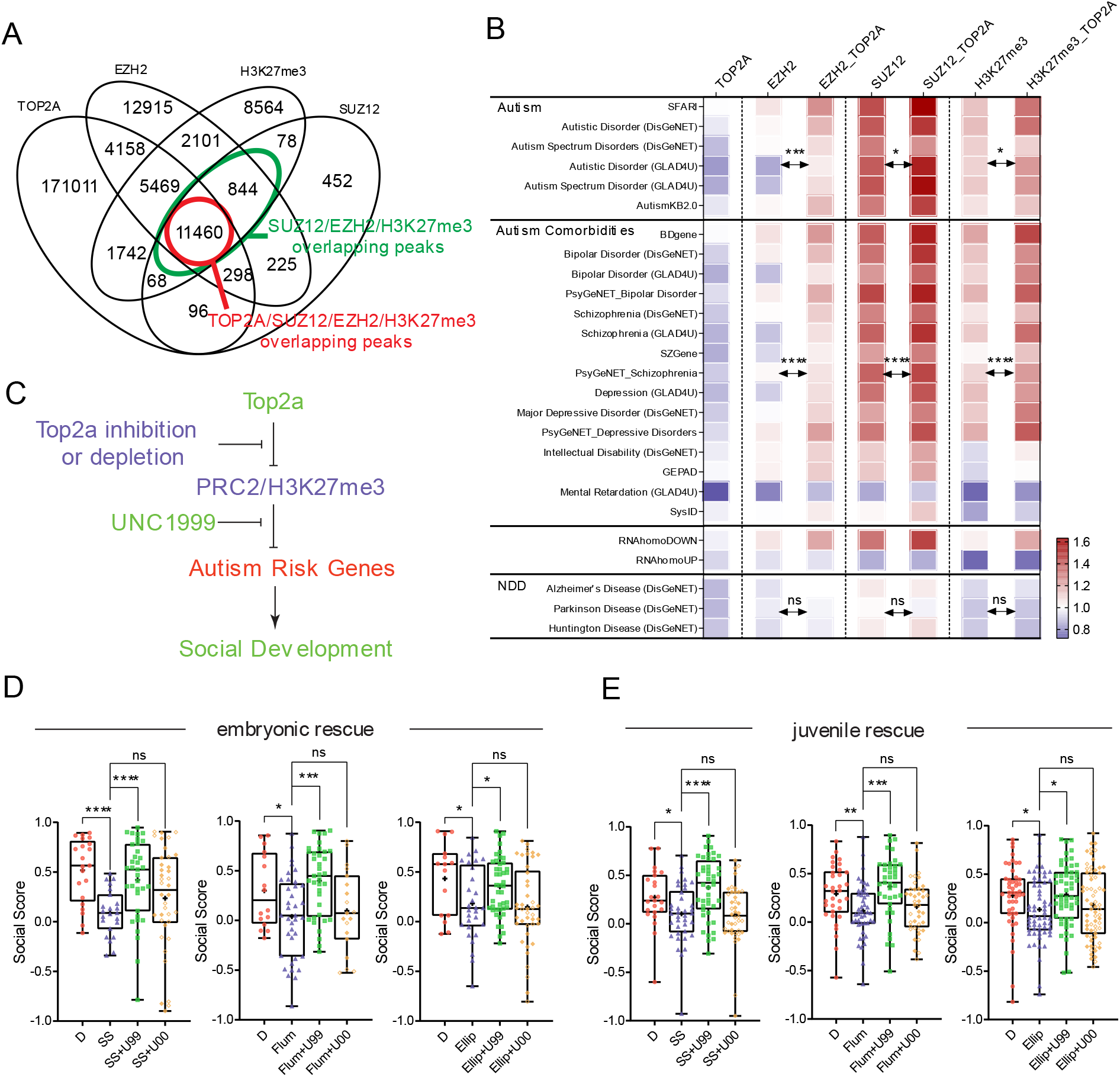
PRC2 and H3K27me3 mediate the Top2a-dependent gene regulation. (**A**) TOP2A binding peaks significantly overlap with EZH2, SUZ12, and H3K27me3. (**B**) Among all EZH2, SUZ12, and H3K27me3 binding peaks, those overlap with TOP2A are more enriched for autism and its comorbidities risk genes and can4Dn, but not for can4Up and neurodegenerative disorder risk gene sets. Significance was calculated by paired t test. NDD: neurodegenerative disorders. (**C**) A working model for how PRC2 and H3K27me3 mediate the Top2a-dependent gene regulation and social development. (**D**) Embryonic (0-3 dpf) co-treatment of 10 μM UNC1999 (U99) with 100 μM sodium salicylate (SS, n=35), 15 μM flumequine (Flum, n=37), and 0.5 μM ellipticine (Ellip, n=44), respectively, rescues social deficits caused by SS (n=23), Flum (n=37), and Ellip (n=29) alone compared to DMSO control (D; left to right: n=21, 16, 15). UNC1999’s inactive analog UNC2400 (U00) does not rescue social deficits (left to right: n=38, 19, 42). (**E**) Social deficits caused by embryonic treatment of SS (n=38), Flum (n=45), and Ellip (n=61) compared to DMSO (left to right: n=23, 34, 47) are rescued by overnight exposure to U99 (left to right: n=44, 39, 56) but not U00 (left to right: n=42, 41, 67) at the juvenile stage (20-21 dpf). ns: not significant, *: *p*<0.05, ***: *p*<0.001, ****: *p*<0.0001.

We assessed the specificity of this similarity by running a blast-style search (UES-blast) against control gene sets from a reference dataset consist of DisGeNET, GLAD4U, and an RNA-seq gene set library RNAseqGEO^49,50,65^ to identify gene sets with the lowest blast scores (highest similarity rankings) to a query gene set (Figure 6B). Using SFARI genes as a query gene set for UES-blast, we found that can4Dn, together with several independently curated autism risk gene sets, rank among the highest, while can4Up ranks much lower. Removing overlapping SFARI genes from can4Dn (can4Dn-SFARI) does not significantly affect its ranking, indicating that the overlapping SFARI genes were not the sole contributors to their rankings. Autism comorbid conditions also rank high on the list, except for intellectual disability (Figures 6C & S10A). We also acquired several neurodegenerative disorder risk gene sets including Alzheimer’s disease, Parkinson’s disease, and Huntington’s disease from DisGeNET as negative controls due to their neurological nature, late onset, and lack of comorbidity with autism; these control gene sets rank relatively low on the list as expected. Using tSNE analysis we found that can4Dn, SFARI, and other autism risk gene sets tightly cluster together, compared to the control disease gene sets in DisGeNET (Figure 6D) or the entire reference dataset (Figure S11). These results suggest that the Top2a-dependent can4Dn and autism risk genes share a very similar and unique set of upstream regulators.

Among the strongly associated upstream regulators, we found 4 that are shared by can4Dn and SFARI genes, but not by can4Up, including SUZ12, EZH2, REST, and SMAD4 (Figure 6B). Intriguingly, SUZ12 and EZH2 have been reported to bind to TOP2A^66^. Because both SUZ12 and EZH2 are core components of the PRC2 complex, a histone methyltransferase, we explored enrichment of histone modifications in our gene sets using ORA analysis and two curated libraries containing published ChIP-seq data gathered from the ENCODE and Roadmap Epigenomics projects^65^. PRC2 methylates histone H3 on lysine 27 (i.e., H3K27me3). Remarkably, we found that can4Dn and SFARI genes, but not can4Up, are highly enriched in genes marked by H3K27me3 but not by the other 29 histone markers examined (Figure 6E). We calculated a H3K27me3 score to quantify this enrichment (Methods). can4Dn and autism risk gene sets give the highest H3K27me3 scores when compared to control gene sets from the reference dataset, while can4Up scores zero as expected (Figures 6F & S10B). Autism comorbid conditions except for intellectual disability also give high H3K27me3 scores.

### PRC2 and H3K27me3 mediate Top2a-dependent gene regulation

We compared genome-wide binding patterns of TOP2A and PRC2 using published ChIP-seq data^63,67^ in human K562 cells, and found that among the triple co-binding sites of SUZ12, EZH2, and H3K27me3, > 93% were also bound by TOP2A (Figure 7A). In addition, while TOP2A binding peaks alone are not enriched for autism risk genes (Figure 7B, first column), overlapping TOP2A binding peaks with EZH2, SUZ12, and H3K27me3 binding peaks, respectively, enriched their selective targeting to autism risk genes and can4Dn, but not to can4Up and the neurodegenerative disorders gene sets (Figure 7B). These results suggest that the selective regulation of autism risk genes by Top2a may be achieved through its interplay with PRC2 and H3K27me3.

While H3K27me3 functions as a gene silencing mark^68^, Top2a binding is strongly associated with open chromatin^63,69^. A FAIRE-Seq analysis revealed that Top2a inhibition by ICRF-193 in mouse embryonic stem cells results in reductions in chromatin accessibility^69^, demonstrating a role for Top2a in maintaining open chromatin. These results suggest an antagonistic relationship between Top2a and H3K27me3. We therefore reasoned that Top2a inhibition may serve to derepress PRC2 on genes contributing to social development, whereas this effect might be mitigated by a PRC2 inhibitor (Figure 7C).

To test this hypothesis, we added an Ezh2 inhibitor UNC1999 and its inactive control compound UNC2400^70^ alongside Top2 inhibitors during embryonic exposure and found that UNC1999 effectively ameliorates social deficits caused by flumequine (a fluoroquinolone antibiotic), ellipticine (a Top2 poison), and sodium salicylate (a Top2a-selective Top2 catalytic inhibitor), while UNC2400 shows no effective rescue (Figure 7D). Perhaps more significantly, we found that overnight exposure to UNC1999 but not UNC2400 the night before Fishbook test effectively rescued social deficits in juvenile fish treated with Top2 inhibitors during embryonic development (Figure 7E). These findings suggest that a significant portion of the social deficit phenotype may be attributed to elevated H3K27me3 that was introduced by Top2a inhibition at the embryonic stage and persisted to the juvenile stage.

## DISCUSSION

Association studies have now identified hundreds of genetic variants that each contribute a small amount of autism risk^7-10^. Given the large number of genes involved and the inability of any one gene to explain more than a small amount of increased risk, one might ask whether the many autism risk genes are regulated independently or are coordinately regulated in some way. Evidence of coordination might point toward shared mechanisms of autism etiology, as well as give hope that autism spectrum disorders of diverse genetic backgrounds might be treatable by targeting their shared mechanisms.

Although genetic factors are clearly important contributors to ∼60% of autism risk, environmental factors appear to explain an additional ∼40% of risk^11,12^. Despite the importance of identifying environmental factors contributing to autism, few have been identified. Interest in identifying environmental risk factors of autism is intense; even a fraudulent claim linking vaccinations and autism prompted lasting changes in vaccine utilization^71^. The difficulty of linking environmental factors with neurobehavioral development may be largely attributed to the massive number of environmental factors and the limited throughput of current research methodologies. Only a fraction of the chemicals in our environment have been examined for potential adverse effects on neurobehavioral development^72,73^. Current experimental methods mainly rely on rodent models. Although these models can often recapitulate complex behavioral phenotypes that relate to human diseases, the time, space, and financial resources required can be prohibitive for conducting large scale experimental screens. Models enabling large scale testing of the impact of small molecules on social development are needed.

The zebrafish possesses an evolutionarily conserved social brain network^15^. Many components of this network have been studied in rodent models for social-related behaviors, such as social reward^74^, aggression^75,76^, mating^77,78^, social learning^79^, and social memory^80^, which paved the way for using zebrafish to investigate social behavior and autism. The Fishbook platform enables systematic evaluation of environmental chemicals to identify those affecting social behavior development. In a systematic small molecule screen, we identified Top2a inhibitors as profound disruptors of social development in zebrafish and mice. Importantly, disruption of Top2a preferentially alters the expression of many of the genes previously associated with autism in humans. Hence, Top2a may serve as a common regulatory factor controlling a large set of genes known to confer autism risk, and as such may also serve as a link between genetic and environmental contributors to autism.

Prior studies have implicated Top2a in autism via different approaches. One study identified *TOP2A* as a member of a core autism risk gene network consisting of 234 genes by analyzing whole exome sequencing data^36^. Another revealed *TOP2A* as one of the 23 hub-genes related to brain maldevelopment in autism toddlers^37^. The Top2 inhibitor ICRF-193 has been shown to inhibit transcription of extremely long genes that are associated with autism in cultured mouse cortical neurons, embryonic stem (ES) cells and ES-cell-derived neurons^35^. In corroboration of both this finding and ours, a FAIRE-Seq analysis found that ICRF-193 caused reduction in chromatin accessibility in ES cells, especially in long silent genes^69^. Interestingly, several Top2 inhibitors were found to reactivate the dormant allele of the autism and Angelman syndrome gene *Ube3a* in primary cortical neurons from mice by reducing transcription of an imprinted antisense RNA *Ube3a-ATS*^34^.

Fluoroquinolones are generally avoided during pregnancy due to their association with developmental toxicity in animals. Despite this consensus, fluoroquinolones are occasionally prescribed to pregnant woman^81,82^, likely due to providers being unaware of the pregnancy or unaware of the potential risk. The results presented here may call for additional investigation into whether prenatal fluoroquinolone exposure in humans increases autism risk. It is noteworthy that a number of known or suspected environmental risk factors of autism are Top2 inhibitors, such as Bisphenol A (BPA)^26,83,84^, polychlorinated biphenyls (PCB)^27,28,85-87^, chlorpyrifos and chlorpyrifos oxon^29,88-90^, and genistein^91-93^. Top2 inhibitors are widely present in the environment, and their possible link to autism has previously been proposed^94^. Chemicals with Top2 inhibiting activities that are present in the environment but have not yet been clearly associated with autism include flavonoids^92,95^, alternariol^96^, Ginkgo biloba leaf extract^97^, and lignans^98-100^, to name a few. Environmental pollution by drugs with Top2 inhibitor activities has also been reported^101-105^. Nevertheless, it is not known whether or not Top2a inhibition is a meaningful contributor to autism prevalence in humans. Further epidemiological studies will be needed.

It is perhaps surprising that Top2a, an enzyme often regarded as performing ‘housekeeping’ functions, might have a specific effect on development of sociality. In fact, as described here, Top2a binds DNA throughout the genome, with no detected preference for genes associated with autism. The situation changes, however, when considering Top2a colocalized with the PRC2. Colocalization of Top2a, PRC2, and the PRC2-deposited epigenetic mark H3K27me3 is highly enriched among autism risk genes, including those discovered through human GWAS studies and those identified in zebrafish in this study. Hence, Top2a appears to influence the expression of diverse genes that together regulate social development by antagonizing PRC2/H3K27me3 mediated gene silencing. In alignment with this hypothesis, mutations in both H3K27 demethylases, KDM6A and KDM6B, have been associated with autism^106,107^, linking defects in a H3K27me3 antagonizing mechanism with autism etiology. Why so many autism risk genes would be co-regulated by the Top2a/PRC2 mechanism is unclear. One possibility is that many genes are required for successful social development, and that being able to simultaneously regulate their expression during development is advantageous.

The effects of Top2a inhibition on sociality appear to be durable: zebrafish and mice treated transiently with Top2a inhibitors during development exhibit social deficits weeks later in the case of zebrafish and months later in the case of mice. This durability may reflect a permanent developmental defect, for example if Top2a inhibition during development alters the structure of the brain in such a way that normal sociality cannot be recovered even after Top2a activity is restored. Alternatively, the social deficits may reflect gene expression changes that are preserved by epigenetic marks but might theoretically be reversed by a second epigenetic intervention. Remarkably, we found that manipulation of the Top2a/PRC2 pathway using the Ezh2 inhibitor UNC1999 could rescue social deficits caused by Top2 inhibition, not only when co-treated during development but even in older animals. In particular, overnight exposure to UNC1999 rescued social deficits caused by embryonic Top2 inhibition, suggesting that a short-term modulation of this pathway at the juvenile stage may be sufficient to boost pro-social gene expression. This phenomenon, if conserved in mammals, may offer hope that Top2a-mediated social deficits might be reversible through epigenetic reprogramming.

In summary, this study reveals a crucial role for Top2a in promoting the development of social behavior in zebrafish and mice. Our results suggest that Top2a is indispensable for maintaining an upstream regulatory network that selectively controls the expression and epigenetic status of a large subset of autism risk genes. Top2a likely functions by antagonizing PRC2 and H3K27me3 mediated gene silencing. Further analyses are warranted to examine potential links between prenatal exposures to Top2-inhibiting chemicals and autism risk in humans through epidemiological and toxicological approaches.

## MATERIALS AND METHODS

### The Fishbook system and Social Score

The basic unit (test arena) of this system is a 3D printed, 10 mm deep, 8.5 mm wide, and 80 mm long rectangular chamber. Each chamber is divided into three compartments by two transparent acrylic windows (1.5 mm thick): a 60 mm long middle testing chamber to place the test subject, and two 8.5 mm long end chambers to place the social stimulus fish or remain empty, respectively. The walls are 1.5 mm thick. The 3D printing model was created using Tinkercad (Autodesk). These arenas were 3D printed using white PLA (polylactic acid) at 100 % infill. Printed arenas were glued onto 3/16” thick white translucent (43% light transmission) acrylic sheets (US Plastic) using a silicone sealer (Marineland). Transparent acrylic windows were precision cut to 8.5 mm × 10 mm pieces using a laser cutter and inserted into printed slots in the arena and fastened using silicone sealer.

The key component of our imaging system is a 322 mm diameter bi-telecentric lens (Opto Engineering) with an IR (850 nm) bandpass filter (Opto Engineering). Videos were taken by a Blackfly 5.0 MP Mono USB3 Vision camera (PointGrey) at 7.5 frames per second (fps). Ambien light was provided using white LEDs (EnvironmentalLights). The arenas were illuminated from below with infrared (850 nm) LEDs (EnvironmentalLights). Infrared LEDs were cooled by a heat sink (H S Marston). Structural supports and enclosure were custom built using parts purchased from Thorlabs, McMaster Carr, and US Plastic.

Test subjects were individually placed inside each test chamber using a plastic transfer pipette with its tip cut off to widen the opening. Their visual access to social stimulus fish were temporarily blocked by a 3D printed white comb-like structure placed in front of the social stimulus compartment (Figures S1E & S1F). Once all test subjects were placed into the test arena array, the array was placed inside the imaging station and the combs were removed to visually expose the social stimulus fish to the test subjects. After a 5 min acclimation period, a 7.5 min (for screening and validation) or 10 min test session was video recorded.

Videos were streamed through the software Bonsai^108^. Videos were analyzed in real-time during recording, and the frame-by-frame x and y coordinates of each fish relative to its own test compartment were exported as a CSV file. Data were analyzed using custom scripts (Python) to calculate social scores and generate tracking plots. Social score was defined as a fish’s average Y-axis position for all frames. We designated the middle of each test chamber as the origin of Y-axis. We then assigned a value of 1 to the end of the chamber close to the social stimulus fish, and a value of −1 to the other end of the chamber which is close to the empty control compartment. Therefore, in this coordinate system, all social scores have values between −1 and 1. A higher social score demonstrates a shorter average distance between a test fish and a social stimulus fish during a test, which suggests a stronger social preference.

### Zebrafish embryonic chemical treatment

For high-throughput screening, fertilized eggs were transferred into 96-well square-well plates at 12 eggs per well. Each well was filled with 500 μl volume E3 medium. E3 medium was supplemented with penicillin/streptomycin to prevent bacterial contamination. Compounds in the Prestwick chemical library (all dissolved in DMSO) were screened at 1:500 dilution for a final concentration of 20 μM. Negative controls were treated with an equal volume of DMSO. Each compound was added to 2 wells. All or 50% of medium and drugs were changed at 1- and 2-day post fertilization (dpf), respectively. Larvae typically hatch between 2-3 dpf. At 3 dpf, all hatched and live but unhatched larvae treated by the same compound were collected and transferred to a petri dish containing fresh E3 medium without penicillin/streptomycin. At 5-7 dpf, larvae from each petri dish were transferred to a separate nursery tank to be raised to 3 weeks of age for Fishbook assay.

For individual compound treatments, each chemical’s general toxicity level was first estimated by measuring overall embryonic lethality after 3 days of treatment (0-3 dpf). Most biologically active small molecules are toxic to zebrafish embryos when applied at high doses and toxicity is often a good benchmark assessment of effective compound exposure. Each compound was first applied to the embryos from 0-3 dpf by dissolving in E3 medium at a wide range of doses, often ranging from 0.1 μM to 1 mM with 10-fold serial dilutions. Two to three additional rounds of assessments were then conducted to narrow down the dose ranges and identify the LD_50_ of each compound. Once toxicity is assessed, compound treatments began at LD_50_ and simultaneously increased and decreased 2-3 times serially by two-fold (at LD_50_×4, LD_50_×2, LD_50_/2, LD_50_/4, etc.). We have also found that some compounds, when applied at a certain dose, showed little to no toxicity at the embryonic stage, but caused massive lethality at a later stage once these drug-treated larvae entered nursery. For these compounds, further adjustments of compound doses were required to identify their new LD_50_ values with nursery survival taken into consideration. Most inhibitors tested effectively induced social deficits when applied at LD_50_. If a compound showed no effect in inducing social deficits at lower doses, at least one dose at > LD_50_ must be successfully tested before a conclusion could be made regarding the compound’s effectiveness. All compounds were dissolved in DMSO (Sigma-Aldrich) or water. The final DMSO concentration was never higher than 1%.

The Peterson lab previously discovered that the logarithm value of a compound’s octanol:water partition coefficient (logP) strongly correlates with the compound’s embryonic permeability^109,110^. A logP value higher than 1 predicts good absorption, whereas compounds with a logP value lower than 1 are often poorly absorbed by the zebrafish embryo. This principle has been confirmed by the author’s own observations, as in our hands, compound with logP values lower than 1 often failed to cause any detectable toxicity even when applied at extremely high doses in E3 medium. This problem can be overcome by injecting compounds directly into embryos at the 1- or 2-cell stage^109^. Therefore, compounds with logP values lower than 1, including ICRF-193, etoposide, bleomycin, and hydroxyurea, were directly injected into embryos. Their respective dosages were determined by first injecting a range of doses, often ranging from 0.1 μM to 10 mM with 10-fold dilutions in between, to assess their general toxicity levels as quantified by the percentage of embryonic lethality induced at each dosage. The doses were then narrowed down and refined to identify the LD_50_ of each compound. Finally, each drug was injected at its LD_50_ dose and at several doses higher and lower (with 2-fold differences) than LD_50_.

### High-throughput screening and validation

A total of 2 rounds of screening and 1 round of validation were performed. The first round exposed 24 embryos to each compound. Treatment groups with social scores lower than 0.1 were identified. These compounds were then hand-picked for the second round of screening, which exposed 48 embryos to each compound. Treatment groups that again inhibited social score to < 0.1 were identified. These compounds were purchased from commercial sources and used to treat 100 embryos. Compounds that significantly reduced social score compared to DMSO control were identified as hits.

### Chemical library and other compounds

The Prestwick library (Prestwick Chemical) contains 1,120 approved drugs dissolved in DMSO at a stock concentration of 10 mM. Other compounds were purchased from the following sources. From Cayman Chemical: flumequine, oxolinic acid, etoposide, mitoxantrone (hydrochloride), ellipticine, bleomycin (sulfate), hydroxyurea, actinomycin D, triptolide, UNC1999. From Sigma-Aldrich: lomefloxacin hydrochloride, ofloxacin, sodium salicylate. From ApexBio: amsacrine. From Santa Cruz: ICRF-193, sobuzoxane, aclarubicin (aclacinomycin A). From Tocris: UNC2400. All individually purchased compounds were dissolved in DMSO or water. Chemical structures were generated using PubChem Sketcher.

### Zebrafish juvenile chemical treatment and Fishbook testing

For the juvenile rescue experiments, 20 dpf zebrafish were collected from nursery tanks. Fish that received the same treatment at the embryonic stage but raised in different tanks were pooled together and then sorted into deep petri dishes (such as a 25 mm deep petri dish) containing 40 ml E3 medium mixed with rotifers as fish feed and 5 μl AmQuel (Kordon) to remove harmful ammonia excreted by fish. 10-15 fish were sorted into each dish. Compounds were then added to each dish. Dishes were incubated at 28 °C in a larvae incubator overnight. On 21 dpf, each dish is fed lightly with rotifers in the morning. Fishbook assay was conducted ∼1-2 hours after feeding. Immediately before plating fish in each petri dish into the Fishbook test arena, the content of the petri dish is poured through a nylon tea strainer so that all liquid passes through and fish are kept in the tea strainer. The tea strainer containing fish is then consecutively dipped into 3 petri dishes containing E3 to wash the residual chemical away from the fish. The fish is then poured into a petri dish containing clean E3 and transferred into the Fishbook test arena using a plastic transfer pipette for testing.

### Site-directed mutagenesis

Construct containing full-length human *TOP2A* cDNA sequence were purchased from Genecopoeia. Site-directed mutagenesis were conducted using QuikChange Site-Directed Mutagenesis Kit (Agilent). The forward and reverse mutagenic primers for *TOP2AY165S* are 5’-aatttggctccagagccatttcgaccacctgtcact −3’ and 5’-agtgacaggtggtcgaaatggctctggagccaaatt-3’.

### Morpholino injection and validation

Splice-blocking morpholinos targeting zebrafish Top2a (5’-GACATTCATCATAAACTCACCCAGA −3’) and Top2b (5’-ATGCAGTAATCTTACCAAGGATCTC −3’) were purchased from Gene Tools. To validate their knockdown efficiencies, injected embryos were collected for mRNA extraction using RNeasy Mini Kit (Qiagen) at 1 dpf, followed by reverse transcription using QuantiTect Reverse Transcription Kit (Qiagen) and PCR using primers that flank the targeted exons. Top2a-MO knockdown efficiency was examined using the forward and reverse primers 5’-GGCTCTCAGTAAGCCCAAGAA −3’ and 5’-TACTGGAGGTCAGGAGTTGGC −3’, with the full-length amplicon predicted to be 401 bp and splice-blocked amplicon 310 bp long. Top2b-MO knockdown efficiency was examined using the forward and reverse primers 5’-AAATGAATGGCCGGGGAGATG −3’ and 5’-TGATGGTTGTCATGTTCTTGTCTCT −3’, with the full-length amplicon predicted to be 297 bp and splice-blocked amplicon 206 bp long. The toxicity of the morpholinos were examined and the doses determined by following a protocol described in the section “Zebrafish embryonic chemical treatment”.

### mRNA overexpression and validation

mRNA *in vitro* syntheses were conducted using the mMESSAGE mMACHINE T7 Transcription Kit (Invitrogen). Linearized constructs containing *TOP2A* and *TOP2AY165S* genes were used as templates. Following synthesis, mRNAs were A-tailed using E. coli Poly(A) Polymerase (NEB) to improve translation efficiency. *TOP2A* and *TOP2AY165S* mRNAs were injected to the yolk of 1-to 2-cell stage embryos. For validation of protein overexpression, ∼200-300 embryos were dechorionated 3-6 hours after injection by 5 min incubation in 1 mg/ml pronase (Roche) in E3 medium in a 2% agarose-coated petri dish and washed with excess amount of fresh E3. Dechorionated embryos were transferred to a 1.5 ml tube and deyolked by pipetting up and down 3 to 5 times using a P200 tip in 200 μl ice-cold PBS. 1 ml ice-cold PBS was added, and the resulting mixture was centrifuged at 300 g for 30 seconds. The supernatant was discarded, and the cell pellet was washed with an additional ml of ice-cold PBS. Cells were lysed in 20 μl RIPA buffer (Santa Cruz Biotechnology) supplemented with cOmplete Mini EDTA-free protease inhibitor tablet (Roche), 5 mM sodium orthovanadate (New England Biolabs), and 10 mM sodium fluoride (Sigma-Aldrich), for 50 min on ice with vortexing every 10 min. The resulting lysate was centrifuged at 14000 g for 15 min at 4°C. The entire volume of supernatant (typically corresponding to ∼30 μg protein) was saved and subsequently combined with 5× Laemmli buffer prior to gel loading for Western blot analysis. Protein concentration was measured by Bradford (Bio-Rad) assay. Anti-TOP2A (Cell Signaling D10G9) and anti-β-actin (Cell Signaling D6A8) antibodies were used for detection. Signal was detected by ECL.

### Zebrafish mutagenesis and genotyping

The complete protocol used to generate and genotype the mutant lines *nrg1*^*-/-*^ and *disc1*^*-/-*^ was previously described^111^. Briefly, frameshift mutations were engineered into the *nrg1* and *disc1* genes of TuAB strain zebrafish at exons 2 and 4 respectively, using CRISPR-Cas9 mediated genome editing. The CRIPSR gRNA spacer sequences were 5’-GGCCGAGGGAGTGGTGCTGG −3’ (*nrg1*) and 5’-GGATACATGCGGTCTGAGCC −3’ (*disc1*). The primers used for genotyping were 5’-TAGGCCGAGGGAGTGGTGCTGG −3’ and 5’-AAACCCAGCACCACTCCCTCGG −3’ for *nrg1*, and 5’-TAGGCTCAGACCGCATGTATCC −3’ and 5’-AAACGGATACATGCGGTCTGAG −3’ for *disc1*. A 20 bp deletion (GGATACATGCGGTCTGAGCC) was detected in exon 2 of *disc1* gene, resulting in a premature stop codon (Figure S1E). A 22 bp deletion (CCAGCACCACTCCCTCGGCCAA) was detected in exon 4 of *nrg1* gene, also resulting in a premature stop codon (Figure S1F).

The mutant line *can4* was genotyped using the forward and reverse primers 5’-CTGCAGAAACCCTGTTAAG - 3’ and 5’-AGGGGATTGACCTCTCGTTG −3’^43^. The mutant line *noto* was genotyped using the forward and reverse primers 5’-TCGGTTCCAAATGTGCTCTCT −3’ and 5’-AGCCTTGCAAGCCCTAATCAT −3’^44^. PCR and subsequent Sanger sequencing were performed using the same primers. Genomic DNA extracted from whole fish following Fishbook assays were purified (Zymo Research) before PCR to remove PCR inhibitors in the fish tissues.

### Mouse prenatal injection

Assessing pregnancy according to the presence of vaginal plugs can be unreliable^112^. We therefore assessed pregnancy based on weight gain following a reported protocol^112^ with modifications. We considered the day after overnight pairwise breeding to be potentially 0 days post coitum (dpc). Females were first weighed on 0 dpc, then single housed until 7 dpc when they were weighed again. Females with weight gains > 1.0 gram were considered pregnant. Drug and vehicle control were injected daily through intraperitoneal injection from 7 dpc to 18 dpc. Births were given on 19 or 20 dpc. ICRF-193 was first dissolved in 100% ethanol (Fisher Scientific), then diluted in saline (Baxter) to a final concentration of 0.01 μg/μl. Diluted drug was injected at a volume of 10 μl per gram body weight to achieve a final dosage of 0.1 mg/kg.

### Social interaction assay

Social interaction was examined as described previously^46^ using 2 months old mice. Briefly, each adult mouse was introduced into a neutral and unfamiliar cage. After a 5 min acclimation period, a foreign age-, sex-, and weight-matched untreated mouse (from separate litters) was introduced into the cage as a social stimulus. Both experimental and untreated females were assayed at the metestrus or diestrus stages to minimize the influence of estrous cycle on social behavior. The test mouse was marked by an odorless black marker at the tail to differentiate from the social stimulus mouse. Testing sessions lasted 5 min and were manually scored for social interactions later. Durations of each social interaction were timed using the software BORIS^113^. All mice were scored blind to their treatment conditions. Social interaction was defined as any period of time in which the test mouse was actively investigating the social stimulus mouse. Investigations included sniffing of the social stimulus mouse toward its facial, abdominal, and anogenital areas, grooming, or closely pursuing the stranger as it explored the cage; investigation of the test mouse by the social stimulus mouse was not scored.

### Novel object investigation assay

The novel object test was performed in a neutral and unfamiliar cage using 8 months old mice in the exact same manner as the social interaction assay, except with a wooden block introduced to the cage after the acclimation period instead of a social stimulus mouse. Total time spent investigating the object was quantified over a 5 min period. Objects were covered with tape during the test; tape was thoroughly removed and replaced in between tests to remove odor traces.

### Ultrasonic vocalization assay

Vocalizations were tested in pups on postnatal days 7 or 8. Cages were acclimatized to the room for 30 min. Pups were individually placed on the test platform in a sound-proof cabinet. Ultrasonic vocalizations were measured for 5 min using an ultrasonic vocalization detector (Avisoft Bioacoustics).

### T-maze assay

T-maze assay was performed as previously described^114^ using 3 months old mice. Each session consisted of eight consecutive trials. In each trial, the mouse was placed in the start compartment of a T-maze. After 1 min before the first trial and 15 sec for the subsequent trials, the door was lifted, and the mouse was left free to explore the two arms of the maze. As soon as the animal entered (with all four paws) one of the two alternative arms (left or right), the door of that compartment was closed for 30 s to confine the animal. The animal was removed between trials and the T-maze was quickly cleaned with 70% alcohol and dried to remove any odor traces that may affect the performance in the next trial.

### Hole-board assay

Hole-board assay was conducted in a 16-hole square apparatus as previously described^46^ using 3 months old mice. Each session lasted 10 min and the total number of head pokes was recorded. The occurrence of perseverative behaviors (manifested as the tendency to explore the same holes) was measured via the coefficient of exploratory variation, calculated as the ratio of the standard deviation of the number of head pokes for each hole over the mean for each mouse. This index^115^ provides a quantitative estimation of the dispersion of the probability distribution of exploratory activity with regard to perseverative behaviors.

### Open field assay

Testing was conducted as described previously^116^ using 10 months old mice. The open field consisted of a grey Plexiglas square arena (40 × 40 cm) enclosed by 4 black walls (40 cm high). On the floor, two zones of equal area were defined: a central square zone of 28.28 cm per side, and a concentric peripheral zone including the area within 11.72 cm from the walls. Mice were placed in the central zone at the beginning of each assay and their behaviors were each monitored for 10 min. Animal motions were tracked using Bonsai. Percentage of time spent in the center zone and average velocity were measured using a custom written Python script.

### Animal husbandry

All animal husbandry and experiment protocols were approved by and carried out in accordance with the Institutional Animal Care and Use Committee at Massachusetts General Hospital or University of Utah.

Fertilized eggs (up to 10,000 embryos per day) were collected from group mating of EkkWill strain zebrafish (*Danio rerio*) (EkkWill Waterlife Resources). Embryos were raised in HEPES (10 mM) buffered E3 medium at 28 °C, with or without compound treatment, during the first 3 days. At 3 days post fertilization (dpf), chorion debris was removed, and larvae were transferred into petri dishes containing fresh E3 medium. At 5-7 dpf, larvae were transferred into nursery tanks and raised at 28 °C on a 14/10 hr on/off light cycle.

Male C57BL/6J mice (*Mus musculus*) were obtained from Jackson Laboratories. Mice were group-housed at no more than 5 per cage and were maintained on a 12/12 hr light/dark cycle. The room temperature was maintained at 20-23 °C with relative humidity at approximately 50%. Food and water were available *ad libitum* for the duration of the study, except during testing. All tests were conducted during the light phase of the light dark cycle.

### Statistical analysis

Graphs were generated using GraphPad Prism or Python. Data were analyzed using 2-tailed Student’s t test. Bonferroni correction was used to adjust for multiple comparison, except for ORA conducted using the enrichR R package. P values less than 0.05 were considered significant.

### RNA-seq and data analysis

We conducted RNA-seq on 3 dpf *can4*^*-/-*^ mutants and their WT siblings. Sequencings and initial data processing were conducted at Huntsman Cancer Institute High-Throughput Genomics and Bioinformatic Analysis Shared Resource. We first incrossed heterozygous *can4* mutants to produce a mixture of *can4*^*+/-*^ (heterozygous), *can4*^*-/-*^ (homozygous), and *can4*^*+/+*^ (wild type; WT) embryos. We then lysed 3 dpf larvae individually by rigorously pipetting in 55 μl DNA/RNA Shield (Zymo Research) at RT followed by overnight incubation at 4°C. We extracted genomic DNA from 5 μl lysate of each larvae using Zymo 96 genomic DNA purification kit and genotyped each sample by PCR and Sanger sequencing. The remaining lysates were stored at −80°C while awaiting genotyping results. Once genotyped, larvae with the same genetic background were pooled for mRNA extraction using Direct-zol RNA Miniprep Kit with TRI Reagent from Zymo Research. We prepared 5 biological replicates for each genetic background – *can4*^*-/-*^ and WT – with each sample containing mRNA extracted from 9 larvae. Libraries were prepared using the Illumina TruSeq Stranded Total RNA Library Prep Ribo-Zero Gold Kit. Sequencing was conducted in NovaSeq with 2 × 150 bp runs at 50 million reads per sample. Sequencing reads were aligned using STAR^117^. Quality control was performed using MultiQC (https://multiqc.info/). Initial data processing and analyses were performed using the R packages DESeq2^118^ and hciR (https://github.com/HuntsmanCancerInstitute/hciR). Data were normalized using the regularized log (rlog) counts method in DESeq2. The *can4*^*-/-*^ mutant data was compared to WT data to identify genes with significantly altered expressions. The raw and processed RNA-seq data have been deposited in the Gene Expression Omnibus under accession number GSE181730.

### Gene sets and gene set libraries

The SFARI^51,52^ gene set was acquired from the archived online SFARI gene database (https://gene-archive-dev.sfari.org/tools/). The AutismKB 2.0^53^ gene set was downloaded from the AutismKB website (http://db.cbi.pku.edu.cn/autismkb_v2/download.php). The BDgene^54^ gene set was downloaded from the BDgene website (http://bdgene.psych.ac.cn/directSearch.do?type=gene&keyword=). The SZGene^56^ gene set was acquired from http://www.szgene.org/.The SysID^57^ gene set was downloaded from SysID database website (https://sysid.cmbi.umcn.nl/table/human-gene-info). PsyGeNET^55^ gene sets for depressive disorders (PsyGeNET Depressive Disorder), bipolar disorder (PsyGeNET Bipolar Disorder), and schizophrenia spectrum and other psychotic disorders (PsyGeNET Schizophrenia) were downloaded from the PsyGeNET website (http://www.psygenet.org/web/PsyGeNET/menu/downloads). The GEPAD (Genomics England’s PanelApp database)^58^ gene set was acquired from the cited publication.

The gene set libraries DisGeNET, RNAseqGEO (RNA-Seq_Disease_Gene_and_Drug_Signatures_from_GEO), ChIP-X (ENCODE_and_ChEA_Consensus_TFs_from_ChIP-X), ENCODE_HM (ENCODE_Histone_Modifications_2015), and Epigenomics_Roadmap_HM (Epigenomics_Roadmap_HM_ChIP-seq) were downloaded from the Enrichr^65^ website (https://amp.pharm.mssm.edu/Enrichr/#stats). The GLAD4U disease gene set library was downloaded from the WebGestalt^119^ website (http://www.webgestalt.org/).

### Over-representation analysis (ORA)

ORA using DisGeNET, KEGG (KEGG_2019_Human), REACTOME (Reactome_2016), and GO term (biological processes, molecular function, and cellular component) libraries (GO_Biological_Process_2018, GO_Molecular_Function_2018, GO_Cellular_Component_2018) were performed in R using the enrichR package (https://cran.r-project.org/web/packages/enrichR/index.html). Results were ranked based on adjusted *p*-value.

Other ORA assays were performed in R using a one-sided Fisher’s exact test with 95% confidence calculated according to the R function fisher.test, and Bonferroni correction was used to adjust for multiple comparison in these tests. Total number of human coding genes was set at 20438 at the time of analysis (GRCh38.p13; https://uswest.ensembl.org/Homo_sapiens/Info/Annotation).

Permutation analysis was conducted by randomly selecting 5000 genes from all 14989 human orthologs of zebrafish genes for each round of permutation. The null distribution consisted of odds ratios calculated for 1000 permutations. The 14989 human orthologs of zebrafish genes were identified using the R package biomaRt^120,121^, based on zebrafish genome assembly GRCz11 and human genome assembly GRCh38.

### Gene set enrichment analysis (GSEA)

GSEA analyses for DisGeNET, GLAD4U, GO, KEGG, and REACTOME libraries were performed using the web-based gene set analysis toolkit^119^ (WebGestalt; http://www.webgestalt.org/). GSEA analysis for the independent disease gene sets were performed using the GSEA software^122,123^.

### Upstream Enrichment Signature (UES) and UES-blast

UES and UES-blast analyses were conducted in R using custom scripts. For each test gene set, we first run an ORA analysis to find significantly enriched upstream regulators using the ChIP-X library (ENCODE_and_ChEA_Consensus_TFs_from_ChIP-X). As suggested by its full name, this library contains gene sets acquired by analyzing ChIP-X data in the ENCODE and ChEA databases. We then calculate a significance score for each significantly enriched (adjusted *p*-value < 0.05) upstream regulator as follows: significance score =- log(adjusted *p*-value). Significance scores for all significantly enriched upstream regulators of a given gene set are normalized to between 0 and 1. All nonsignificant upstream regulators are assigned a significance score of 0. With a score assigned to each upstream regulator, we created a signature for each gene set which we named Upstream Enrichment Signature (UES).

We constructed a reference dataset containing >1700 UESs from analyzing published disease risk genes including DisGeNET and GLAD4U, and an RNA-seq gene set library RNAseqGEO. To avoid variations caused by gene sets containing too few genes, only gene sets containing ≥500 genes from the DisGeNET library and ≥250 genes from the GLAD4U library were analyzed. We then conduct an unbiased search using a custom-written blast-style querying algorithm (UES-blast) to find gene sets with the lowest blast scores (highest similarity rankings) to a query gene set, e.g., SFARI genes. UES-blast was conducted by first calculating the Euclidean distances between test gene sets and the target gene set (e.g., SFARI genes), and then ranking test gene sets based on their distances to the target gene set.

### Histone modifications enrichment analysis and H3K27me3 score

We assessed enrichment of histone marks in the promoter regions of genes in each test gene set using ORA analyses. For each given histone mark, the curated ENCODE_HM and Epigenomics_Roadmap_HM libraries often contain a number of query gene sets each listing genes that are targeted by this histone mark in their promoter regions in a specific cell or tissue type. For example, a query gene set named “H3K27me3 cerebellum mm9” contains all genes that are marked by H3K27me3 in the mouse cerebellum, based on analyzing published ChIP-X data. In our analysis, we run ORA analyses for each test gene set against all query gene sets in the two libraries to identify those that are enriched in our test gene set. We then count the number of significantly enriched query gene sets for each histone mark, such as the data shown in Figure 7E.

To specifically quantify enrichment of H3K27me3 in each test gene set, we calculated a H3K27me3 score as follows: H3K27me3 score = the ratio of significant H3K27me3 gene sets over all H3K27me3 gene sets in the library × the ratio of significant H3K27me3 gene sets over all significant histone mark gene sets = (number of significant H3K27me3 gene sets / number of all H3K27me3 gene sets) × (number of significant H3K27me3 gene sets / number of all significant histone marks gene sets). As an example, if a test gene set is significantly enriched for 60 H3K27me3 gene sets out of a total number of 100 H3K27me3 gene sets in the library, and at the same time also significantly enriched for another 10 assorted histone marks gene sets, then its H3K27me3 score is calculated as (60 / 100) × (60 / (60 + 10)) = 0.6 × 6/7 = 0.51. For each test gene set, its final H3K27me3 score was acquired by first calculating two separate H3K27me3 scores using the ENCODE_HM and Epigenomics_Roadmap_HM libraries, respectively, and then combining these two scores by summation. All analyses were conducted in R using custom scripts.

### ChIP-seq data analyses

Genome-wide TOP2A binding data was acquired from GEO accession number GSE79593^63^; we used the DMSO-treated sample in this dataset for analyses. EZH2, SUZ12, and H3K27me3 ChIP-seq data were acquired from GEO accession numbers GSM1003576, GSM1003545, and GSM733658, respectively. To enable meaningful comparison with TOP2A binding peaks, all ChIP-seq data were acquired from the same cell line K562 cells. Peaks from the EZH2, SUZ12, and H3K27me3 that overlap with TOP2A peaks were identified using the R package GenomicRanges^124^. Genes that are targeted by the original peaks or the overlapping peaks at their promoter or gene body regions were mapped using the R package bumphunter^125,126^ and exported for subsequent analyses. Venn diagram for peak overlaps was drawn using the R package ChIPseeker^127^.

## ACKNOWLEDGEMENTS

The *can4* and *noto* zebrafish mutants were generously provided by Dr. Leonard Zon’s lab and Dr. Herwig Baier’s lab, respectively. We thank members of our research groups, in particular A. Rennekamp, Y. Jin, L. Mosher, G. Floris, and R. Cadeddu, for helpful advice. We thank the University of Utah Comparative Medicine Center, the Centralized Zebrafish Animal Resource, and the Massachusetts General Hospital Cardiovascular Research Center zebrafish facility for animal husbandry. We also thank the University of Utah Huntsman Cancer Institute High-Throughput Genomics and Bioinformatic Analysis Shared Resource for genomics experiments and initial analyses.

## Funding

This work was supported by the L. S. Skaggs Presidential Endowed Chair and by the National Institute of Environmental Health Sciences of the National Institutes of Health under Award Number K99ES031050. The content is solely the responsibility of the authors and does not necessarily represent the official views of the National Institutes of Health.

## Author contributions

Y.G. conceived the study, built the equipment, designed and conducted the experiments, wrote the codes, analyzed the data, and wrote the manuscript; S.C.G. designed the rodent experiments; A.K.N. and J.J.Y. designed the CRISPR knockout experiments; T.Z., B.R.P., D.L.H., and I.A.D. conducted the experiments; M.B. designed the rodent experiments, interpreted the data, and edited the manuscript; R.T.P conceived the study, designed the experiments, interpreted the data, and wrote the manuscript. All authors contributed meaningful insights during discussions and reviewed and approved the final version of the manuscript.

## Competing interests

The authors declare no competing interests.

## Data and materials availability

The raw and processed RNA-seq data have been deposited in the Gene Expression Omnibus under accession number GSE181730. All other data are available in the main text or supplementary materials. Code is available on the GitHub repository: https://github.com/yijie-geng/Fishbook and are archived on Zenodo under DOI 10.5281/zenodo.5171582. Reagents are available from the corresponding author upon reasonable request.

## Notes

https://www.ncbi.nlm.nih.gov/geo/query/acc.cgi?acc=GSE181730

https://github.com/yijie-geng/Fishbook

https://zenodo.org/record/5171582#.YUjLm7hKiUk

### Competing Interest Statement

The authors have declared no competing interest.

