## Supplementary Materials for "Top2a promotes the development of social behavior via PRC2 and H3K27me3"

**This PDF file includes:**

Figs. S1 to S11
Movies S1
Data S1

**Other Supplementary Materials for this manuscript include the following:**

Movies S1
Data S1

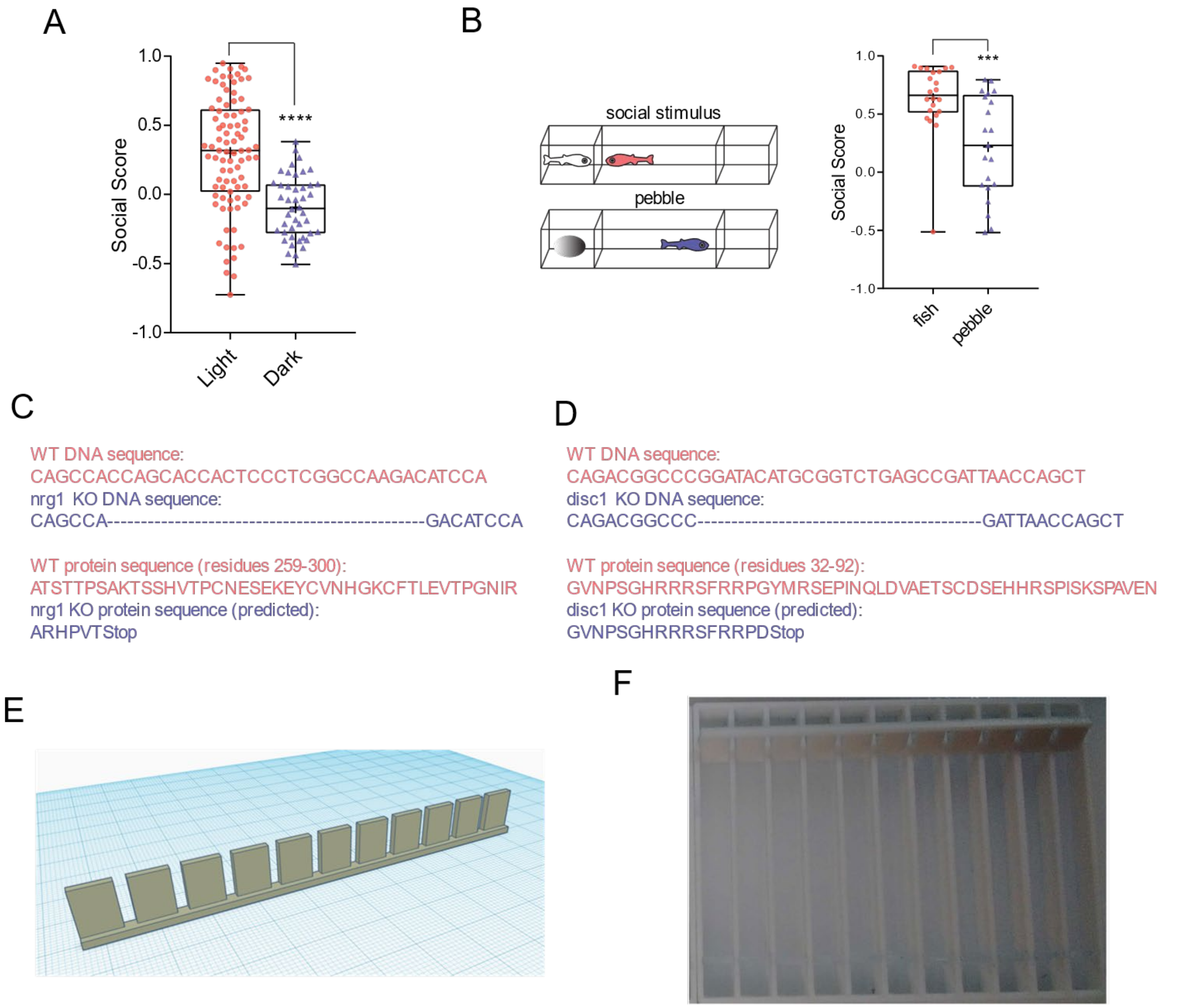


**Figure S1.** (**A**) Boxplot comparing social scores of WT fish in light (n = 87) or dark (n = 44). ****: $p < 0.0001$. (**B**) Boxplot comparing social scores of WT fish in response to a social stimulus (fish; n = 22) or pebble (n = 21). ***: $p < 0.001$. (**C**) DNA and protein sequences of WT and *nrg1* knockout fish. (**D**) DNA and protein sequences of WT and *disc1* knockout fish. (**E**) 3D printing design for the view-blocking comb. (**F**) Image of a comb placed on a Fishbook test arena. The comb is not inserted in this image.

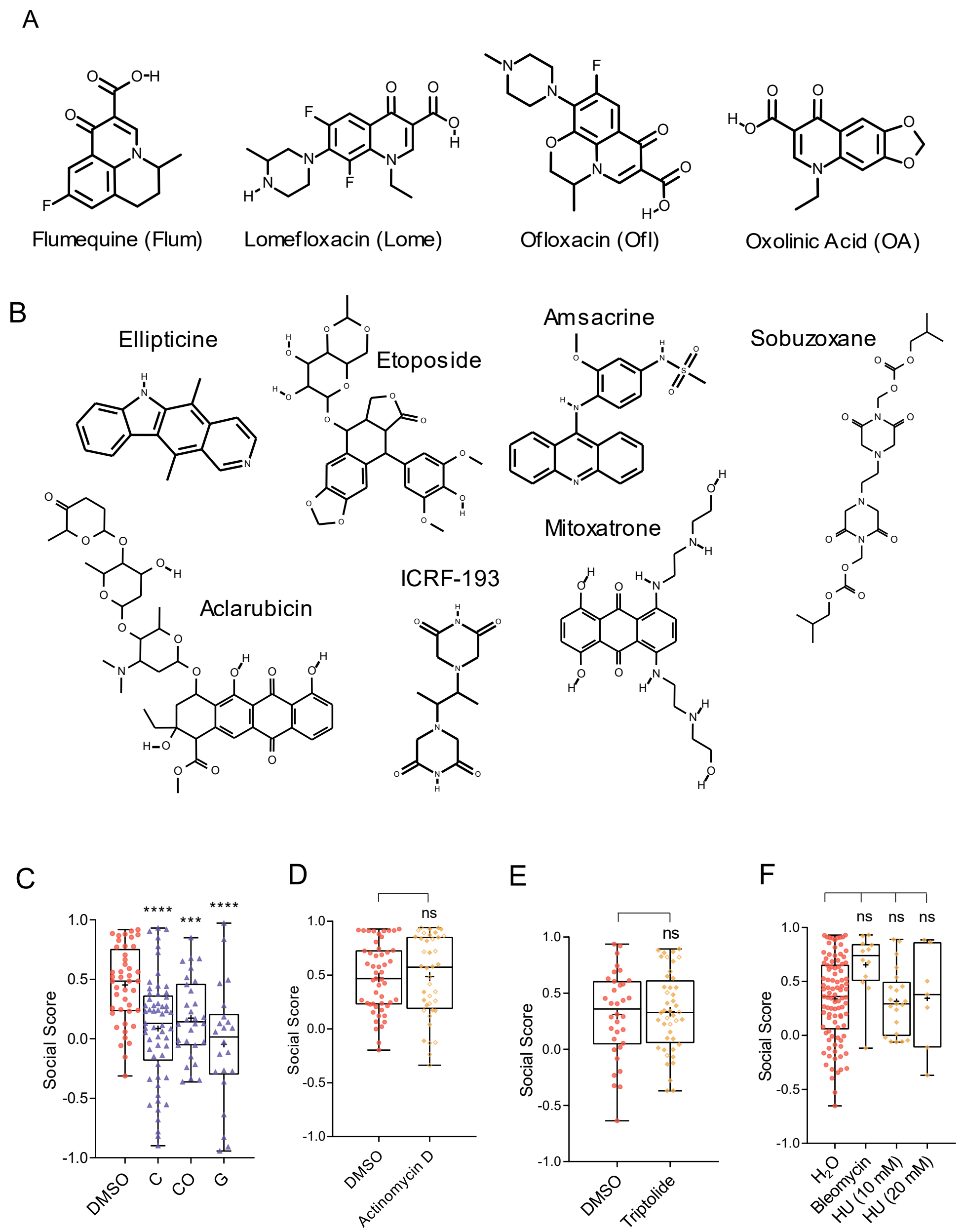


**Figure S2.** (**A**) Chemical structures of fluoroquinolones. (**B**) Chemical structures of eukaryotic Top2 inhibitors. (**C**) Boxplot showing social scores of fish treated with DMSO (n=44), chlorpyrifos (C; 5 µM; n=57), chlorpyrifos oxon (CO; 1 µM; n=30), and genistein (G; 8 µM; n=22). (**D**) Boxplot comparing social scores of fish treated with DMSO (n = 51) or actinomycin D (20 µM; n = 40). (**E**) Boxplot comparing social scores of fish treated with DMSO (n = 34) or triptolide (0.2 µM; n = 44). (**F**) Boxplot comparing social scores of fish treated with DMSO (n = 97), bleomycin (1 µM; n = 12), or hydroxyurea (HU; 10 mM, n = 21; 20 mM, n = 7).
ns: not significant, ***: p<0.001, ****: p<0.0001.

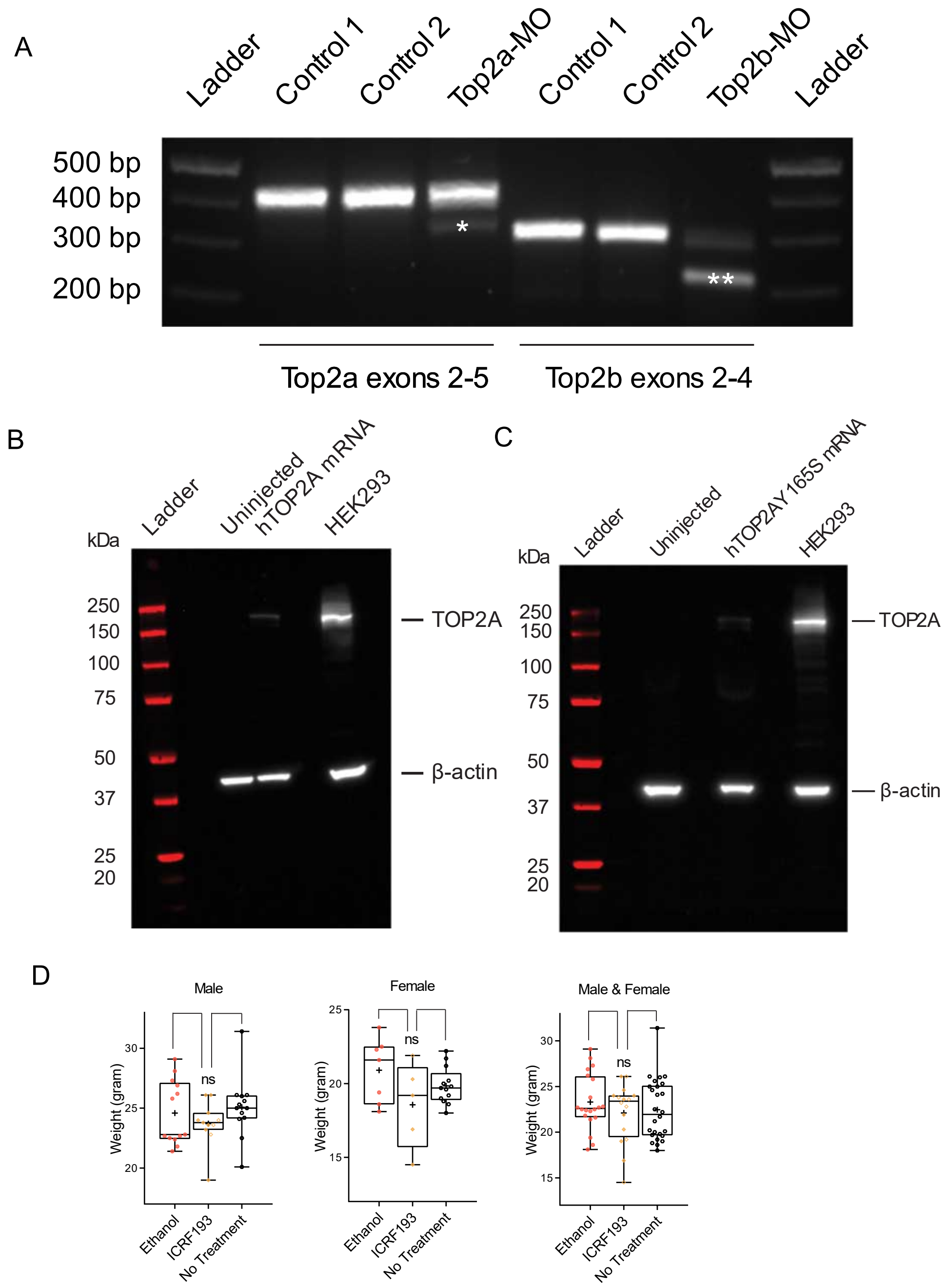


**Figure S3.** (**A**) DNA gel image showing RT-PCR result of 1 dpf embryos injected with Top2a-MO (0.05 mM), Top2b-MO (0.05 mM), or un-injected (Controls 1 & 2). n = ~ 80 – 100 embryos for each condition. *: splice-blocked amplicon of Top2a. **: splice-blocked amplicon of Top2b. (**B, C**) Western blot images showing protein overexpression following injection of 250 ng/µl *hTOP2A* (B) and *hTOP2AY165S* (C) mRNAs in zebrafish embryos. HEK293 cell lysate was used as positive controls. Uninjected zebrafish embryos were used as negative controls. β-actin was used as a loading control. (**D**) Boxplot comparing the body weights of mice treated with ethanol (male: n = 13, female: n = 7), ICRF193 (male: n = 11, female: n = 5), and no-treatment control (male: n = 13, female: n = 13) at 2 months of age. ns: not significant.

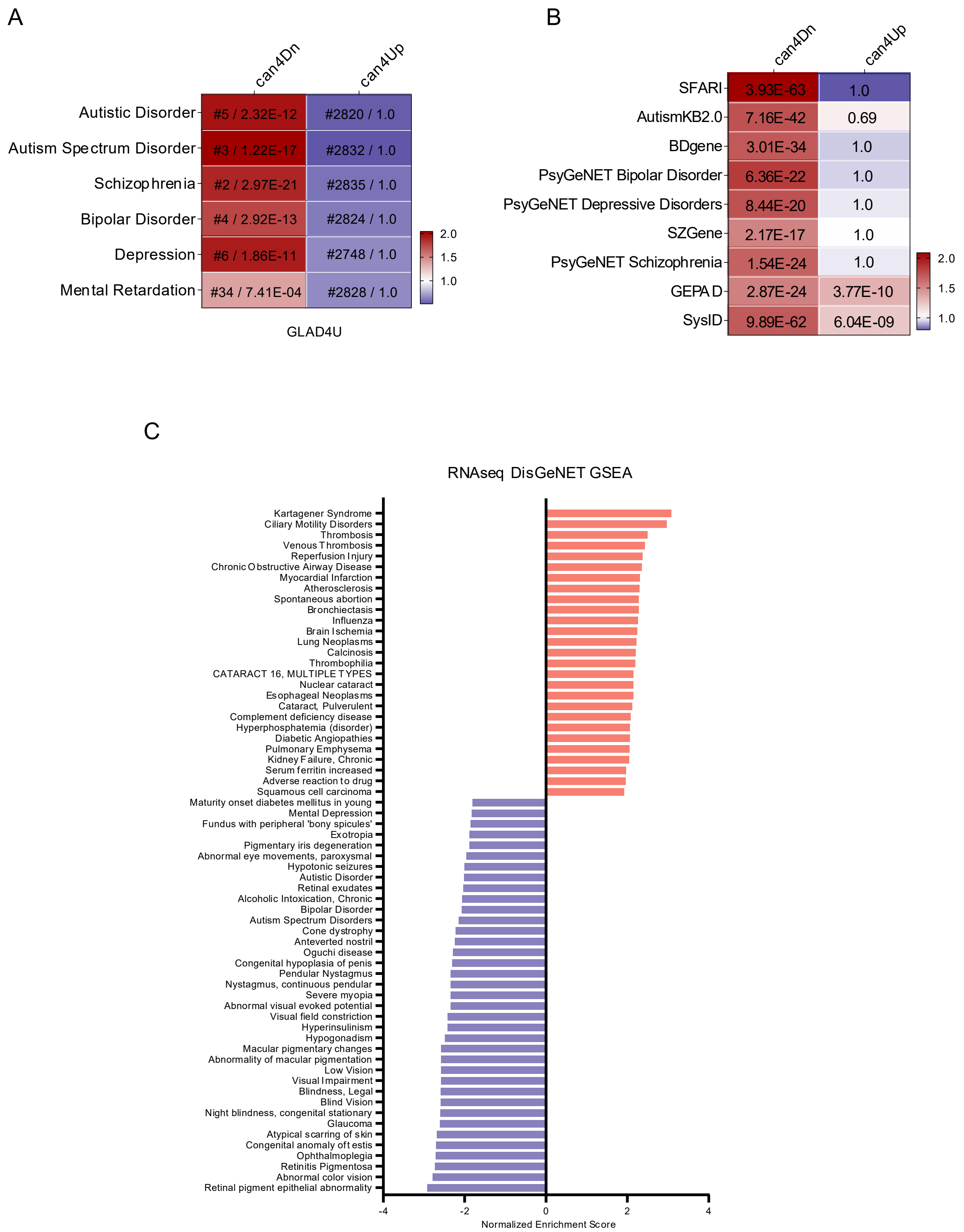


**Figure S4.** (**A**) can4Dn but not can4Up is selectively enriched for autism and its comorbidities risk gene sets from the GLAD4U library. For each cell, color represents odds ratio, and numbers represent ranking out of 3071 diseases in GLAD4U library (before slash) and adjusted *p*-value (after slash). (**B**) Heatmap showing ORA analysis comparing can4Dn and can4Up using several independent disease risk gene sets related to autism and its comorbid disorders. Color in each cell represents odds ratio values. Number in each cell represents adjusted *p*-value. (**C**) Bar chart showing normalized enrichment scores from GSEA analysis for RNA-seq data using the DisGeNET library.

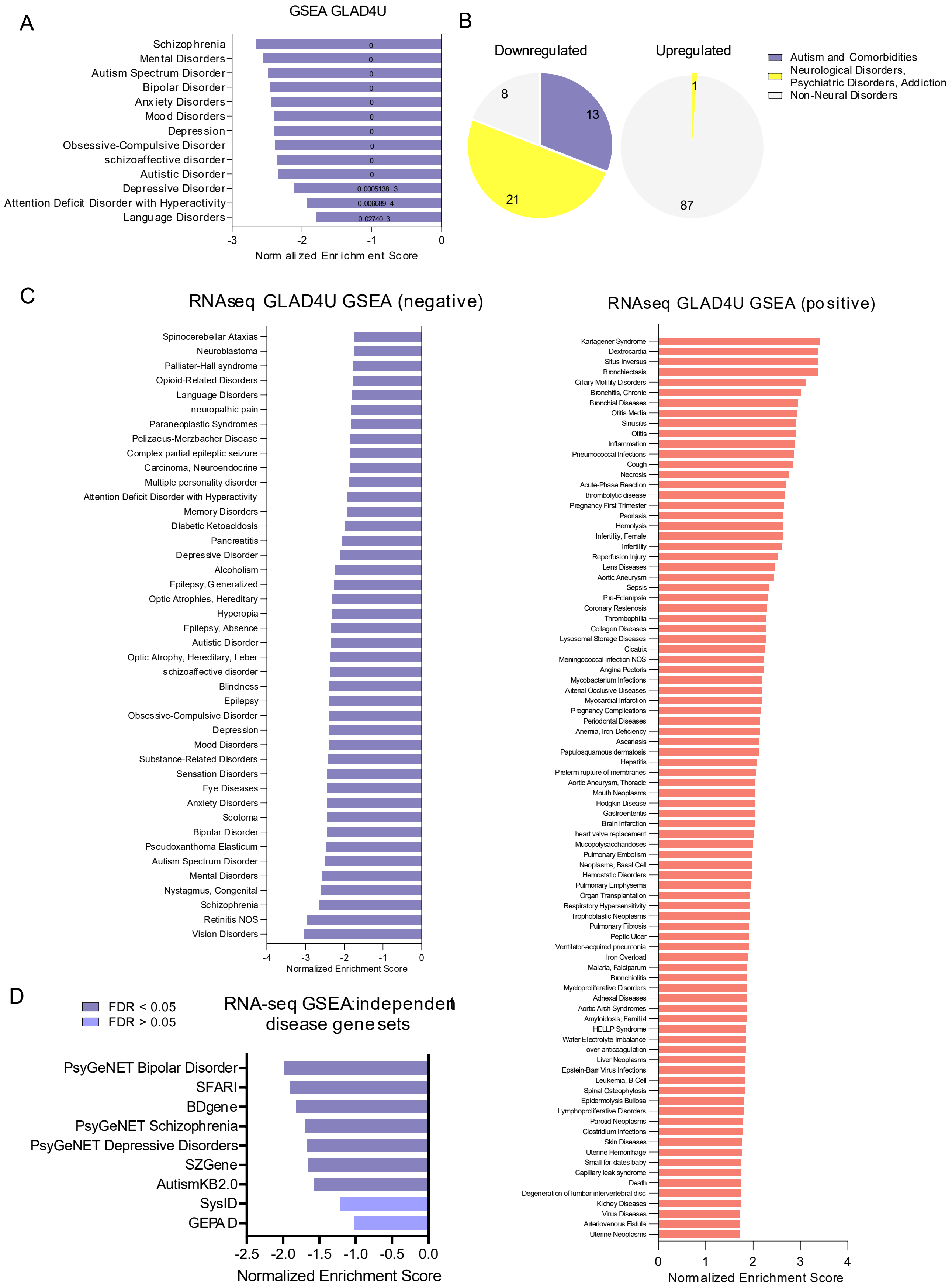
A
GSEA GLAD4U
Schizophrenia
Mental Disorders
Autism Spectrum Disorder
Bipolar Disorder
Anxiety Disorders
Mood Disorders
Depression
Obsessive-Compulsive Disorder
schizoaffective disorder
Autistic Disorder
Depressive Disorder
Attention Deficit Disorder with Hyperactivity
Language Disorders
0
0
0
0
0
0
0
0
0
0
0.0005138 3
0.006689 4
0.02740 3
-3
-2
-1
0
Normalized Enrichment Score
B
Downregulated
Upregulated
8
13
21
1
87
Autism and Comorbidities
Neurological Disorders,
Psychiatric Disorders, Addiction
Non-Neural Disorders
C
RNAseq GLAD4U GSEA (negative)
Spinocerebellar Ataxias
Neuroblastoma
Pallister-Hall syndrome
Opioid-Related Disorders
Language Disorders
neuropathic pain
Paraneoplastic Syndromes
Pelizaeus-Merzbacher Disease
Complex partial epileptic seizure
Carcinoma, Neuroendocrine
Multiple personality disorder
Attention Deficit Disorder with Hyperactivity
Memory Disorders
Diabetic Ketoacidosis
Pancreatitis
Depressive Disorder
Alcoholism
Epilepsy, Generalized
Optic Atrophies, Hereditary
Hyperopia
Epilepsy, Absence
Autistic Disorder
Optic Atrophy, Hereditary, Leber
schizoaffective disorder
Blindness
Epilepsy
Obsessive-Compulsive Disorder
Depression
Mood Disorders
Substance-Related Disorders
Sensation Disorders
Eye Diseases
Anxiety Disorders
Scotoma
Bipolar Disorder
Pseudoxanthoma Elasticum
Autism Spectrum Disorder
Mental Disorders
Nystagmus, Congenital
Schizophrenia
Retinitis NOS
Vision Disorders
-4
-3
-2
-1
0
Normalized Enrichment Score
RNAseq GLAD4U GSEA (positive)
Kartagener Syndrome
Dextrocardia
Situs Inversus
Bronchiectasis
Ciliary Motility Disorders
Bronchitis, Chronic
Bronchial Diseases
Otitis Media
Sinusitis
Otitis
Inflammation
Pneumococcal Infections
Cough
Necrosis
Acute-Phase Reaction
thrombolytic disease
Pregnancy First Trimester
Psoriasis
Hemolysis
Infertility, Female
Infertility
Reperfusion Injury
Lens Diseases
Aortic Aneurysm
Sepsis
Pre-Eclampsia
Coronary Restenosis
Thrombophilia
Collagen Diseases
Lysosomal Storage Diseases
Cicatrix
Meningococcal infection NOS
Angina Pectoris
Mycobacterium Infections
Arterial Occlusive Diseases
Myocardial Infarction
Pregnancy Complications
Periodontal Diseases
Anemia, Iron-Deficiency
Ascariasis
Papulosquamous dermatosis
Hepatitis
Preterm rupture of membranes
Aortic Aneurysm, Thoracic
Mouth Neoplasms
Hodgkin Disease
Gastroenteritis
Brain Infarction
heart valve replacement
Mucopolysaccharidoses
Pulmonary Embolism
Neoplasms, Basal Cell
Hemostatic Disorders
Pulmonary Emphysema
Organ Transplantation
Respiratory Hypersensitivity
Trophoblastic Neoplasms
Pulmonary Fibrosis
Peptic Ulcer
Ventilator-acquired pneumonia
Iron Overload
Malaria, Falciparum
Bronchiolitis
Myeloproliferative Disorders
Adnexal Diseases
Aortic Arch Syndromes
Amyloidosis, Familial
HELLP Syndrome
Water-Electrolyte Imbalance
over-anticoagulation
Liver Neoplasms
Epstein-Barr Virus Infections
Leukemia, B-Cell
Spinal Osteophytosis
Epidermolysis Bullosa
Lymphoproliferative Disorders
Parotid Neoplasms
Clostridium Infections
Skin Diseases
Uterine Hemorrhage
Small-for-dates baby
Capillary leak syndrome
Death
Degeneration of lumbar intervertebral disc
Kidney Diseases
Virus Diseases
Arteriovenous Fistula
Uterine Neoplasms
0
1
2
3
4
Normalized Enrichment Score
D
FDR < 0.05
FDR > 0.05
RNA-seq GSEA: independent
disease gene sets
PsyGeNET Bipolar Disorder
SFARI
BDgene
PsyGeNET Schizophrenia
PsyGeNET Depressive Disorders
SZGene
AutismKB2.0
SysID
GEPAD
-2.5
-2.0
-1.5
-1.0
-0.5
0.0
Normalized Enrichment Score

**Figure S5.** (**A, B**) GSEA analysis of RNA-seq data using the GLAD4U library shows enrichment for autism and its comorbidities risk genes (A & B) and neurological conditions risk genes (B) in downregulated but not upregulated genes. Value inside each bar represents FDR (A). Significance: FDR<5%. (**C**) Bar charts showing normalized enrichment scores (NESs) from GSEA analysis for RNA-seq data using the GLAD4U library. Left: hits with positive NESs; right: hits with negative NESs. (**D**) Bar chart showing normalized enrichment scores from GSEA analysis for RNA-seq data using several independent disease gene sets related to autism and its comorbid disorders. Significance: FDR < 5%.

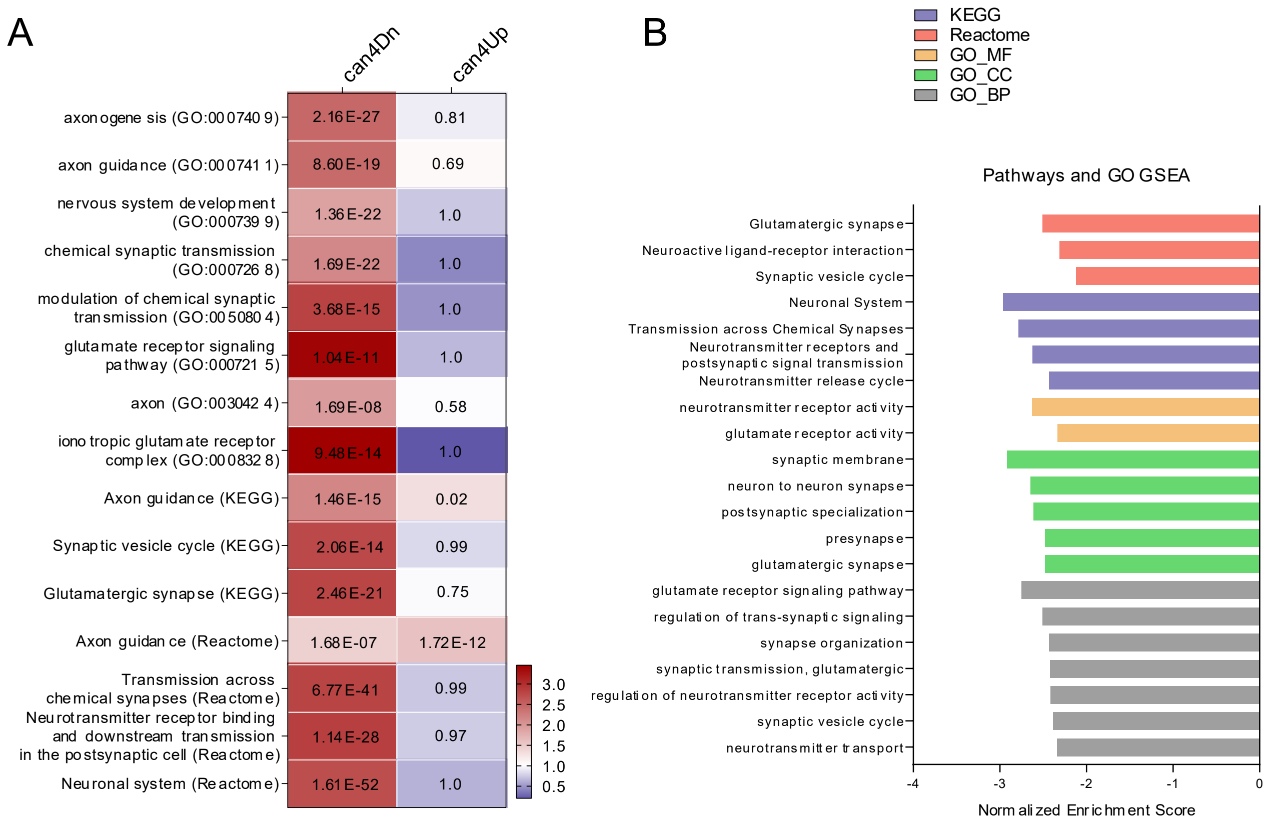


**Figure S6.** (**A**) Heatmap showing ORA analysis comparing can4Dn and can4Up using KEGG, REACTOME, and GO libraries. Color in each cell represents odds ratio. (**B**) Bar chart showing GSEA analysis of KEGG, Reactome, and GO libraries using the RNA-seq data.

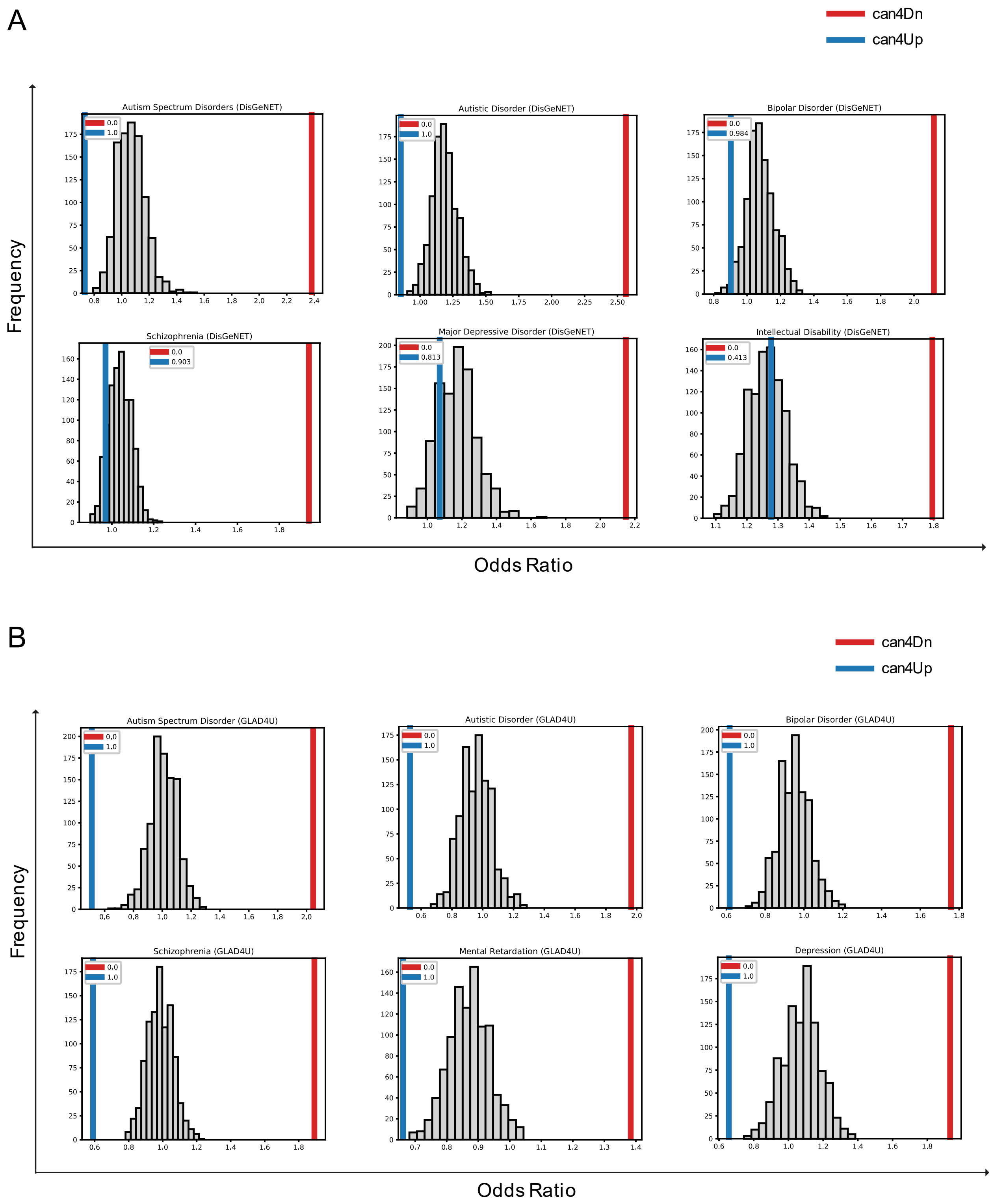


**Figure S7.** (**A**) Histogram showing the null distribution of odds ratios generated by 1000 permutations by randomly selecting 5000 genes out of all human orthologs of zebrafish genes and conduct ORA analysis using selected gene sets from the DisGeNET library. The odds ratios for can4Dn (red) and can4Up (blue) are marked by colored vertical lines, and their respective *p*-values are shown as numbers in legends. A *p*-value <0.008 was considered significant to correct for multiple comparisons. (**B**) Histogram showing the null distribution of odds ratios generated by 1000 permutations by randomly selecting 5000 genes out of all human orthologs of zebrafish genes and conduct ORA analysis using selected gene sets from the GLAD4U library. The odds ratios for can4Dn (red) and can4Up (blue) are marked by colored vertical lines, and their respective *p*-values are shown as numbers in legends. A *p*-value <0.008 was considered significant to correct for multiple comparisons.

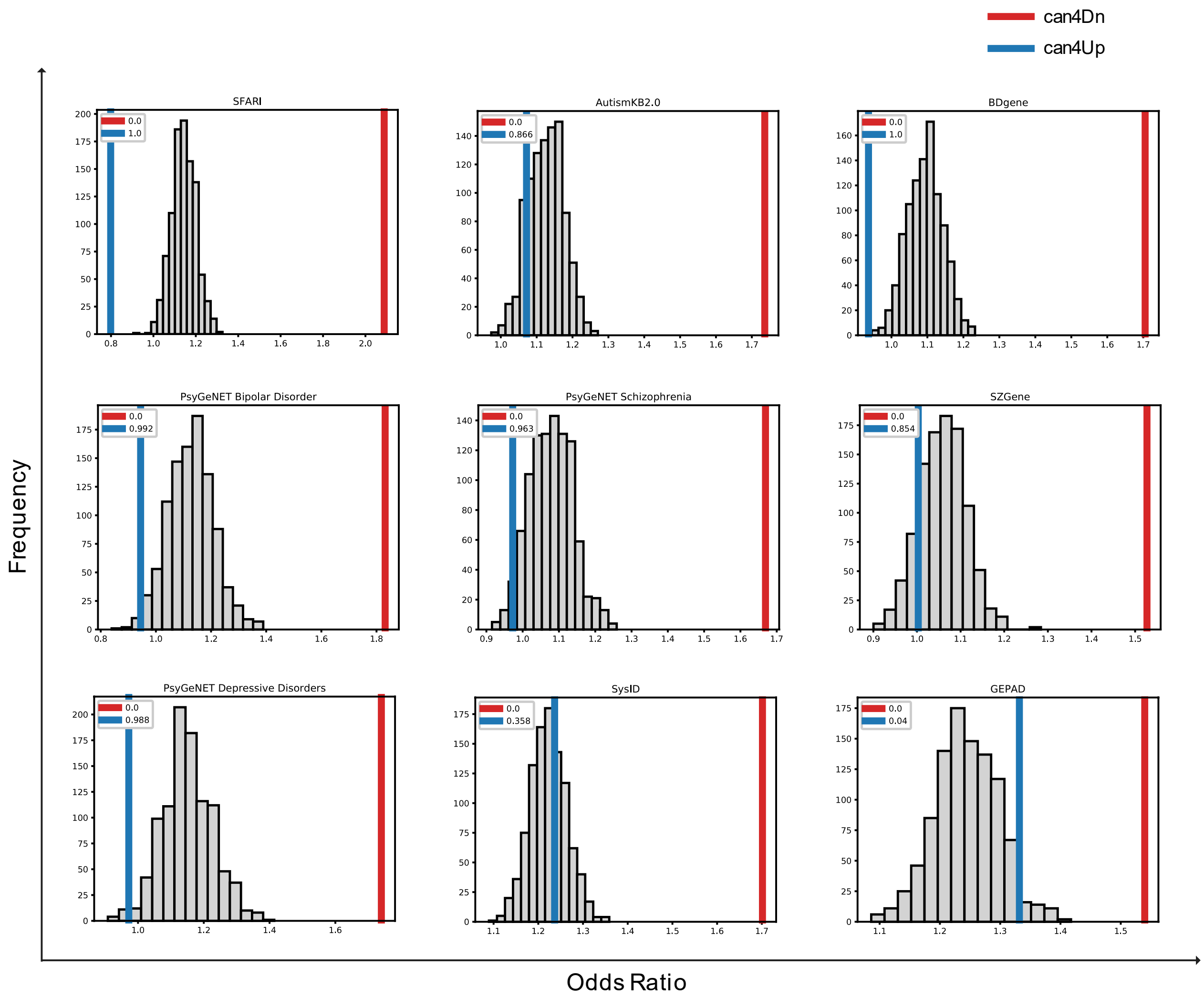


**Figure S8.** Histogram showing the null distribution of odds ratios generated by 1000 permutations by randomly selecting 5000 genes out of all human orthologs of zebrafish genes and conduct ORA analysis using selected gene sets from several independent disease gene sets related to autism and its comorbid conditions. The odds ratios for can4Dn (red) and can4Up (blue) are marked by colored vertical lines, and their respective *p*-values are shown as numbers in legends. A *p*-value <0.005 was considered significant to correct for multiple comparisons.

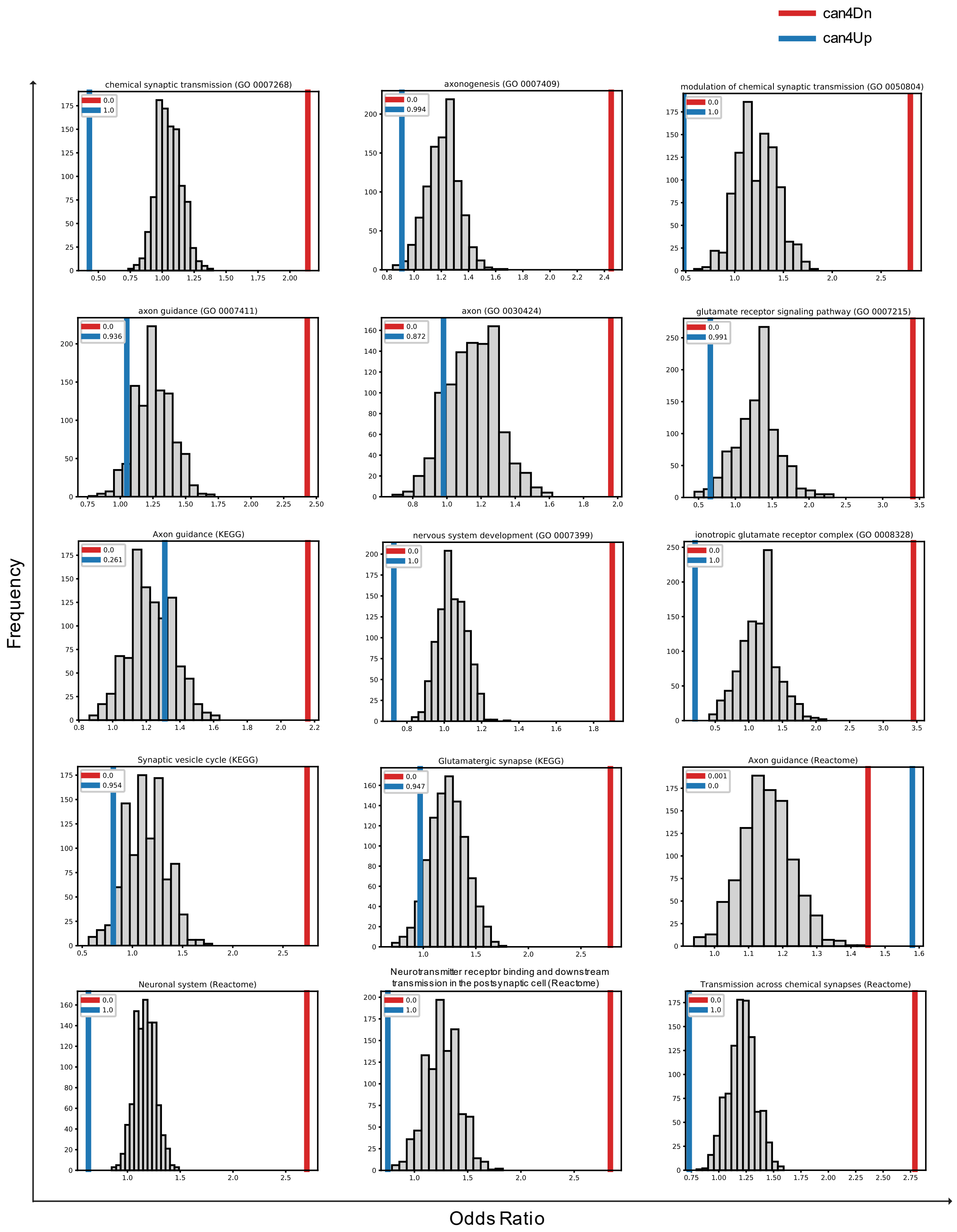


**Figure S9.** Histogram showing the null distribution of odds ratios generated by 1000 permutations by randomly selecting 5000 genes out of all human orthologs of zebrafish genes and conduct ORA analysis using selected gene sets from KEGG, Reactome, and GO libraries. The odds ratios for can4Dn (red) and can4Up (blue) are marked by colored vertical lines, and their respective *p*-values are shown as numbers in legends. A *p*-value <0.003 was considered significant to correct for multiple comparisons.

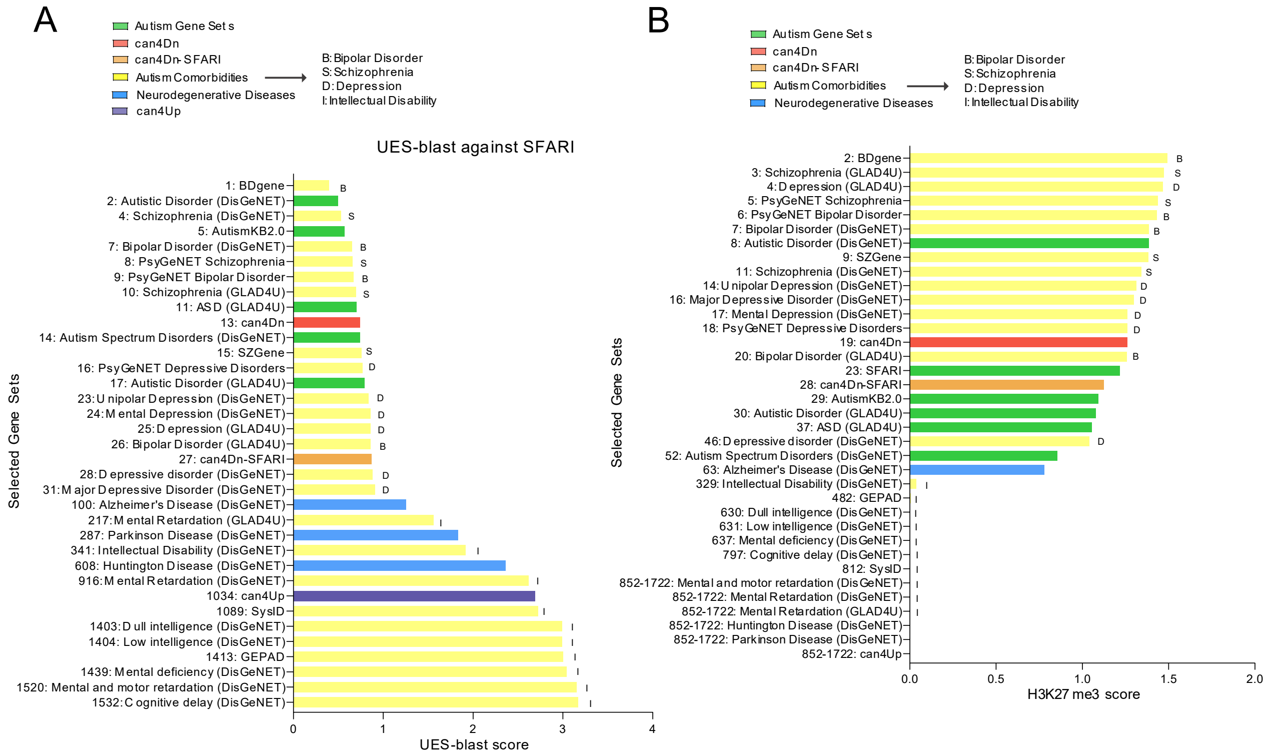


**Figure S10.** (**A**) A detailed look at UES-blast rankings of the key gene sets in Figure 6C. Bar chart shows the UES-blast scores of autism risk gene sets (green), can4Dn (red), can4Dn-SFARI (orange), autism comorbid conditions risk gene sets (yellow; D: depression, B: dipolar disorder, S: schizophrenia, I: intellectual disability), can4Up (dark blue), and neurodegenerative disorders risk gene sets (blue). Numbers in the x-axis labels before gene set names represent the UES-blast rankings of each labeled gene set. (**B**) A detailed look at H3K27me3 score rankings of the key gene sets in Figure 6F. Bar chart shows the H3K27me3 scores of autism risk gene sets (green), can4Dn (red), can4Dn -SFARI (orange), autism comorbid conditions risk gene sets (yellow; D: depression, B: dipolar disorder, S: schizophrenia, I: intellectual disability), can4Up (dark blue), and neurodegenerative disorders risk gene sets (blue). Numbers in the x-axis labels before gene set names represent the H3K27me3 score rankings of each labeled gene set.

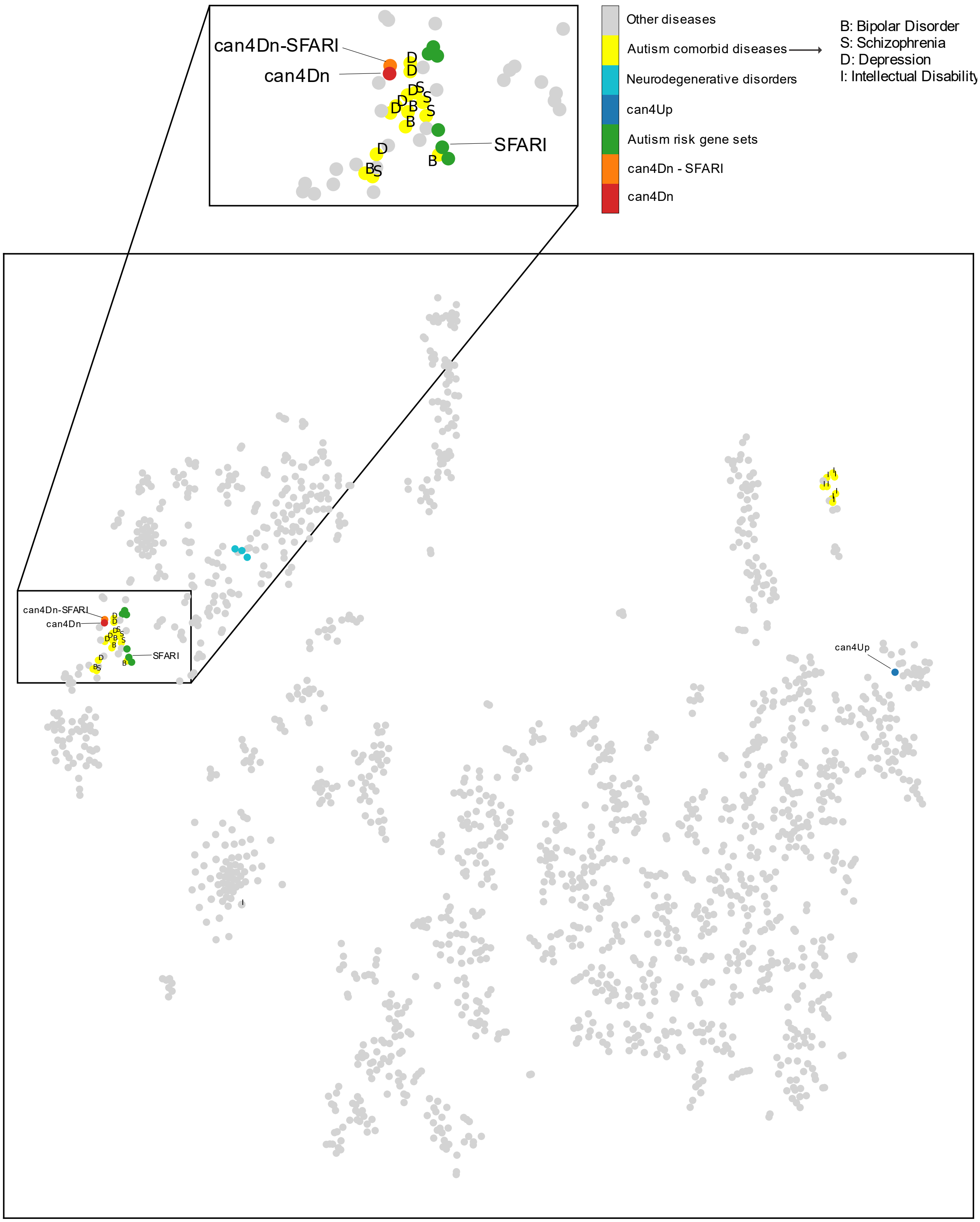


**Figure S11.** tSNE clustering of can4Dn, can4Dn-SFARI, autism risk gene sets, neurodegenerative disorders risk gene sets, and all control gene sets from the reference dataset. Letters label autism comorbidities risk gene sets, I: intellectual disability; D: depression; S: schizophrenia; B: bipolar disorder.

**Movies S1.**
Video recording of a Fishbook test. Played at 4× the actual speed.

**Data S1.**
The can4Dn and can4Up gene lists.
